# Paving the way for synthetic C1- metabolism in *Pseudomonas putida* through the reductive glycine pathway

**DOI:** 10.1101/2022.07.10.499465

**Authors:** Lyon Bruinsma, Sebastian Wenk, Nico J. Claassens, Vitor A.P. Martins dos Santos

## Abstract

One-carbon (C1) compounds such as methanol, formate, and CO_2_ are alternative, sustainable microbial feedstocks for the biobased production of chemicals and fuels. In this study, we engineered the carbon metabolism of the industrially important bacterium *Pseudomonas putida* to assimilate these three substrates through the reductive glycine pathway. First, we demonstrated the functionality of the C1-assimilation module by coupling the growth of auxotrophic strains to formate assimilation. Next, we extended the module from formate to methanol using both NAD and PQQ – dependent methanol dehydrogenases. Finally, we demonstrated CO_2_-dependent growth through CO_2_ reduction to formate by the native formate dehydrogenase, which required short-term evolution to rebalance the cellular NADH/NAD^+^ ratio. This research paves the way to engineer *P. putida* towards growth on formate, methanol, and CO_2_ as sole feedstocks, thereby substantially expanding its potential as a sustainable and versatile cell factory.

## INTRODUCTION

Current biotechnological production of chemicals and fuels primarily depends on sugars and other plant biomass fractions as substrates. However, there are serious sustainability concerns related to these substrates due to their competition for land with biodiversity and food production (Wendisch et al., 2016). Therefore, alternative microbial feedstocks need to be urgently considered. One-carbon (C1) feedstocks are considered prime sustainable alternatives, as CO_2_ or reduced C1-feedstocks can be obtained from abundantly available atmospheric CO_2_ or waste gas streams. CO_2_ can be converted into chemicals and fuels by chemolithoautotrophic microbes when supplied with an inorganic energy source or a reduced C1-source. Moreover, (electro)catalytic methods are increasingly available to efficiently convert CO_2_ into the soluble, reduced C1-molecules formate and methanol (Cotton et al., 2020; Liu et al., 2020; Stöckl et al., 2022). Both were identified as promising microbial feedstocks, given their easy transport, storage, and feeding into the bioproduction process (Claassens et al., 2018; Cotton et al., 2020; Naik et al., 2010; Wendisch et al., 2016).

A major challenge in realizing C1-based bioproduction is that natural C1-utilizers are generally limited in their biotechnological applicability. They are hard to genetically modify, grow slowly, harbor inefficient C1-assimilation pathways, and/or have a narrow product spectrum (Antoniewicz, 2019; Bennett et al., 2018). Attractive biotechnological production organisms, such as *Escherichia coli, Saccharomyces cerevisiae,* and *Pseudomonas putida* are naturally unable to grow on C1-substrates. Nevertheless, in recent years efforts have been made in the former two organisms to establish synthetic methanol and formate assimilation. Especially efforts in *E. coli* have been successful, in which full synthetic methylotrophy and formatotrophy have been established (Chen et al., 2020; Kim et al., 2020)

However, despite the model status of *E. coli*, this bacterium is not optimal for many industrial processes, due to its lack of robustness under full-scale operation. Therefore, in recent years, *P. putida* has emerged as a promising host, as this soil bacterium is naturally endowed to withstand harsh conditions and physiochemical stresses. These features make it an industrially relevant microbe for which the generation of a plethora of products has been demonstrated (Ankenbauer et al., 2020; Martin-Pascual et al., 2021; Nikel & de Lorenzo, 2018; Poblete-Castro et al., 2012) *P. putida* cannot naturally grow on C1-substrates, and so far only the use of formate as an auxiliary energy source has been demonstrated (Zobel et al., 2017). Hence, to realize sustainable bioproduction using C1-substrates in *P. putida*, efficient assimilation pathways must be established.

Several pathways can be considered to establish C1-assimilation in this bacterium. Typical natural pathways for C1-assimilation such as the Calvin Cycle, the Ribulose Monophosphate Pathway (RuMP), and the Serine Cycle can allow for growth on CO_2_, methanol, and/or formate (Antoniewicz, 2019; H. Yu & Liao, 2018). However, all share the disadvantage of having a cyclic architecture and intensive overlap with the host’s native metabolism (Bar-Even et al., 2013). Additionally, the Calvin Cycle and the Serine Cycle are relatively energy-inefficient due to their high ATP consumption. In recent years, the reductive glycine pathway (rGlyP) has been suggested as an alternative pathway for C1-assimilation in model microbes (Claassens, 2021). This ATP-efficient, linear pathway was first designed as a synthetic pathway and recently found in nature as a CO_2_ and formate assimilation pathway (Bar-Even et al., 2013; Figueroa et al., 2018; Sánchez-Andrea et al., 2020). In the rGlyP, formate is converted to 5,10 - methylene - THF through consecutive THF-ligation and reduction reactions. Next, 5,10 - methylene - THF, together with CO_2_, ammonia, and NADH produces glycine in the glycine cleavage system (GCS), which can operate in the reverse direction under high CO_2_ concentrations. Then, glycine will condense with a second molecule of 5,10 - methylene - THF to produce serine (Figure 1).

**Figure 1.**
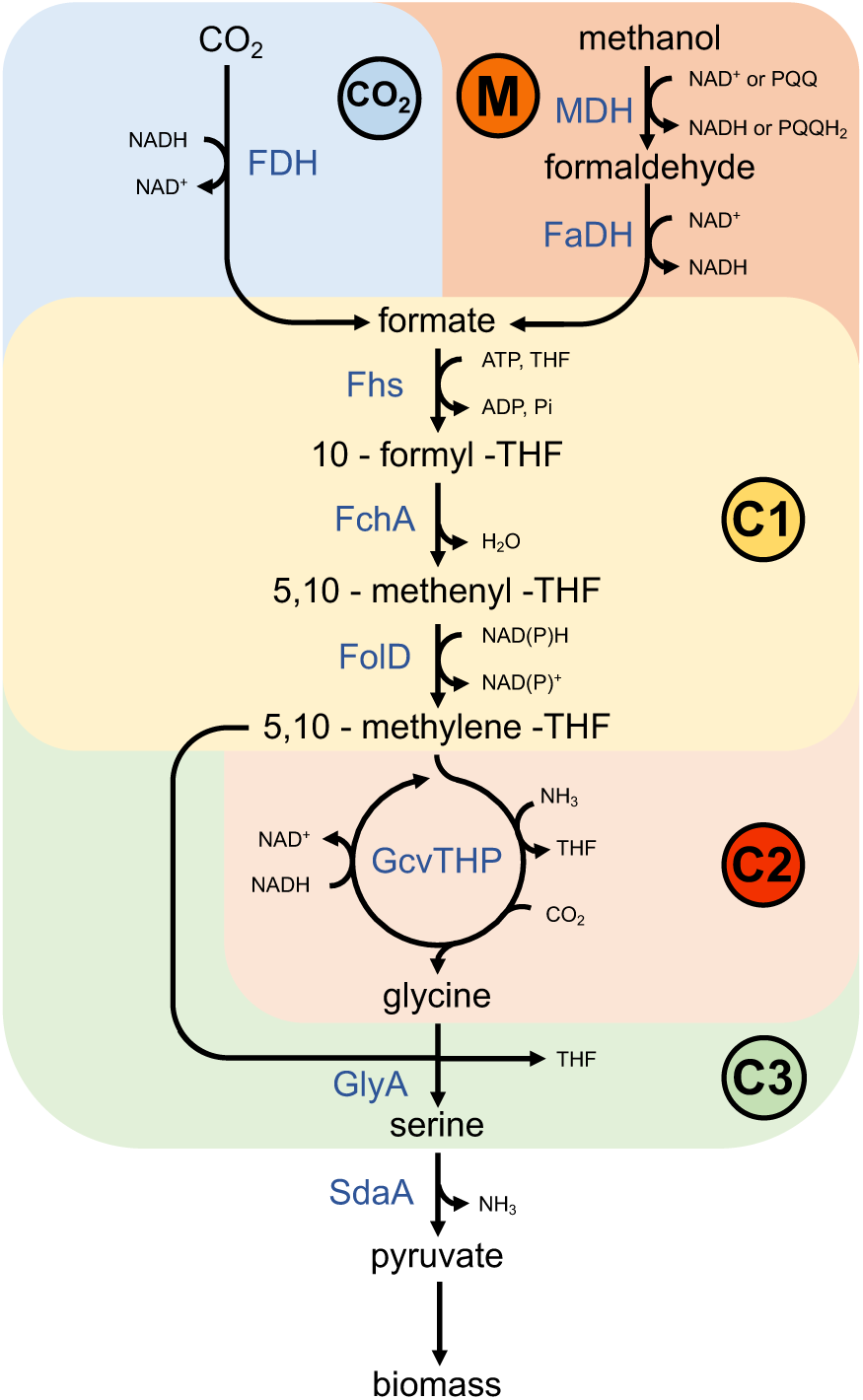
Core module of the reductive glycine pathway implemented in this study. The pathway is divided into several modules. The CO_2_ and M modules convert CO_2_ and methanol to formate, respectively. Formate is then converted to 5,10 methylene-THF in the C1 module. Subsequently, the 5,10 – methylene - THF is converted to glycine by the C2 module comprising the reverse glycine cleavage system. In the C3 module, glycine is condensed with 5,10 methylene-THF to produce serine. Then, serine is deaminated to pyruvate where it provides the cell with the needed biomass. Abbreviations: (FDH), formate dehydrogenase, (MDH), methanol dehydrogenase, (FaDH), formaldehyde dehydrogenase, (Fhs), formate-THF-ligase, (FchA), formyltetrahydrofolate cyclohydrolase (FolD), bifunctional methylenetetrahydrofolate dehydrogenase/methyltetrahydrofolate cyclohydrolase, (GcvTHP), glycine cleavage system, (GlyA), serine hydroxymethyltransferase, (SdaA), serine deaminase, (THF), tetrahydrofolate.

At last, serine is converted to pyruvate through deamination and from thereon to biomass (Bar-Even et al., 2013; Claassens et al., 2022; Yishai et al., 2017, 2018). This formate assimilation pathway can be further extended towards methanol by adding a module containing a methanol and a formaldehyde dehydrogenase. Moreover, when equipped with a CO_2_-reducing formate dehydrogenase (FDH) and an additional energy source, the rGlyP could allow full growth on CO_2_ as the sole substrate (Figure 1).

Modular implementation of the rGlyP has recently led to the full establishment of this pathway in *E. coli* for growth on formate and methanol, and in *Cupriavidus necator* for growth on formate (Claassens et al., 2020; Kim et al., 2020). Another recent work has demonstrated the establishment of the core of the rGlyP in *S. cerevisiae*, by converting formate into glycine (Gonzalez De La Cruz et al., 2019). The assimilation of CO_2_ into the rGlyP has been proposed before, but it has not yet been demonstrated experimentally in an engineered strain (Cotton et al., 2018).

In this study, we establish the foundation for C1-assimilation through the rGlyP for formate, methanol, and CO_2_ in *P. putida*. We follow a growth-coupled modular engineering approach to establish the core of the rGlyP (Orsi et al., 2021). We demonstrate the assimilation of formate, as well as methanol, through the core of the rGlyP into serine. Moreover, we demonstrate, for the first time, a growth-coupled selection for CO_2_ fixation through reverse FDH activity into the rGlyP. Overall, this work demonstrates C1-assimilation of three highly promising C1-feedstocks in *P. putida.* The establishment of complete C1-metabolism will substantially strengthen the position of *P. putida* as an industrial chassis in the bio-industry and thereby contribute to the transitions to a biobased economy.

## RESULTS

### Implementing the reductive glycine pathway

To implement the rGlyP, we split the pathway into three modules: C1, C2, and C3 (Figure 1). The C1 module converts formate to 5,10 - methylene - THF, which is further converted to glycine via the reverse operation of the GCS in the C2 module. In the C3 module, glycine is condensed with another 5,10 - methylene - THF to serine. Finally, serine can be converted to pyruvate and from there to biomass. We constructed two auxotrophic strains in which the functionality of the modules could be coupled to growth, i.e., if the modules are functional the auxotrophy is relieved and the strains will grow. To test the C1 and C3 modules, we constructed a growth coupled design termed C1-S-Aux by deleting the genes of both GCSs (Δ*gcvTPH-I / II*) and the D-3-phosphoglycerate dehydrogenases (Δ*serA /* Δ*PP_2533*) (Figure 2A). This strain is unable to produce serine and the C1-precursor molecules ( e.g. 5,10 - methylene - THF) which are essential for the biosynthesis of purines, thymidine, coenzyme A and methionine (Yishai et al., 2017). In this strain, growth on a canonical carbon source (e.g., glucose) can only be restored when serine is supplemented, or when both C1 and C3 modules are present and active with glycine and formate as substrates. In this case, the C1 module will convert formate into 5,10 - methylene – THF, and the C3 module will condense 5,10 - methylene - THF with glycine to form serine (Figure 2a). To realize the heterologous expression of the C1 module, we created the pC1 plasmid by placing the formate-THF ligase (*fhs*), 5,10 - methenyl-THF cyclohydrolase (*fchA*), and the bifunctional 5,10 - methenyl-THF cyclohydrolase/ 5,10 - methylene - THF dehydrogenase (*fold*) genes from *Clostridium ljunghdahlii* DSM13528 on a SEVAb24 backbone. We equipped the C1-S-Aux strain with pC1 and tested the growth of the strain on a medium containing glucose, glycine, and formate. We observed that the C1 module could carry enough flux into the C1-pool and serine to restore growth of C1-S-Aux. Growth occurred at a similar growth rate (doubling time ∼1 hour) and even a higher biomass yield than for the control medium with glucose and serine (Figure 2b). Overexpression of the C3 module was not necessary as endogenous activity was enough to restore growth. When formate was omitted from the medium, no growth was observed, which reflects its dependency on the assimilation of formate via the C1-module of the rGlyP. To confirm this dependency, we performed labeling experiments with ^13^C-labeled formate and measured the labeling in the proteinogenic amino acids glycine, serine, and histidine (Figure 2c). If the C1 and C3 modules are functioning as expected, serine is derived from the condensation of unlabeled glycine with once-labeled 5,10 - methylene - THF (derived from ^13^C-formate). Histidine biosynthesis requires the incorporation of 10 - formyl - THF, coming from formate in the C1 module, so it should also be labeled once. As expected, both serine and histidine were almost completely labeled once, confirming the activity of the C1 and C3 modules.

**Figure 2.**
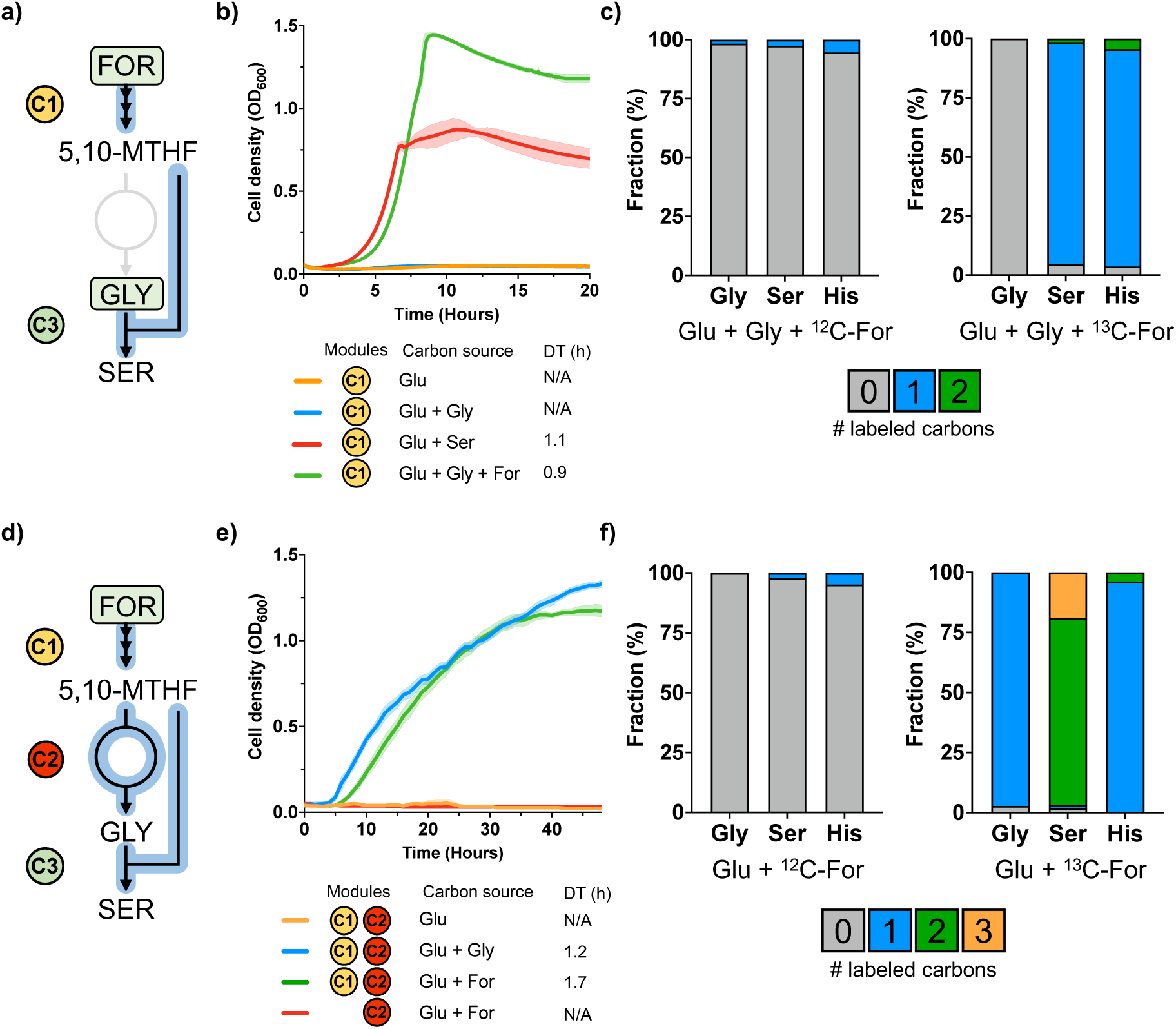
Formate-dependent growth. **a)** Selection of the C1 module in C1-S-aux. Growth is only possible when serine is supplemented to the medium or if both glycine and formate are present. **b)** Growth of C1-S-aux. Overexpression of the C1 module converts formate to replenish the cellular C1- moieties. The C3 module subsequently produces serine, restoring growth. **c)** ^13^C labeling experiments confirm that cellular C1-moieties are produced exclusively from formate. **d)** Selection of the combined activity of the C1, C2, and C3 modules in C1-G-S-Aux. **e)** Growth of C1-G-S-aux. Growth can solely be restored when glycine or formate are added to the medium (at elevated CO_2_ levels through the addition of 100 mM sodium bicarbonate). **f)** ^13^C labeling experiments confirm that cellular C1-moieties, glycine, and serine are produced exclusively from formate and CO_2_. Abbreviations: (Gly) glycine, (Ser), serine, (His), histidine, (For), formate, (5,10-MTHF), 5,10-methylene-THF, (Glu), Glucose. Growth curves and labeling experiments are averages from biological triplicates.

Next, we aimed to test the C2 module, which comprises the reverse operation of the GCS. To test this module, we built a growth-coupled selection strain termed C1-G-S-Aux by deleting the genes encoding threonine aldolase (Δ*ltaE*), and isocitrate lyase (Δ*aceA*), as well as *serA* and PP_2533. This strain is unable to produce serine, glycine, and the C1-precursor molecules and requires external supplementation of serine or glycine for growth on glucose By the combined activity of the C1, C2, and C3 modules, formate together with CO_2_ can generate glycine and subsequently serine to relief the auxotrophy (Figure. 2d). As the native expression level of the GCS is likely not high enough to sustain enough flux, we designed a plasmid to overexpress the native GCS system. We constructed plasmid pC2 (SEVAb65backbone) overexpressing the endogenous *gcvT-I*, *gcvP-I,* and *gcvH-I* genes. We transformed C1-G-S- Aux with the plasmids pC1 and pC2 and were able to restore growth upon the addition of formate, at a slightly lower growth rate (doubling time 1.7 hours) and yield than for the glycine supplemented control. (Figure 2e). Similar to C1-S-Aux, endogenous activity of the C3 module was sufficient to restore growth. Isotopic labeling of the proteinogenic amino acids glycine, serine, and histidine after growth on ^13^C-formate further confirmed the combined activity of the rGlyP modules (Figure 2f). Glycine is produced by the condensation of labeled 5,10 - methylene - THF and unlabeled CO_2_ in the GCS and is expected to be labeled once. Almost all the glycine was labeled once, confirming the combined activity of the C1 and C2 modules. Serine is expected to be labeled twice as its produced from once labeled glycine plus a labeled 5,10 - methylene - THF molecule. As expected, serine was predominantly labeled twice.

### Extending the reductive glycine pathway with methanol assimilation

After demonstrating formate assimilation via the C1, C2, and C3 modules of the rGlyP we wanted to test if these modules could also serve to support methanol assimilation in *P. putida* (Figure 1). Like formate, methanol is a soluble microbial feedstock that can be efficiently generated from CO_2_ and renewable electricity. Methanol is converted to formate in two enzymatic steps. First, methanol is oxidized to formaldehyde by a methanol dehydrogenase (MDH). Second, formaldehyde is further oxidized to formate by a formaldehyde dehydrogenase. From here on formate can be further assimilated through the engineered modules described before. To engineer methanol utilization in *P. putida,* we constructed the M – module comprising the necessary enzymes to oxidize methanol to formate. The genome of *P. putida* encodes for more than 20 alcohol dehydrogenases, yet none of them is annotated as an MDH. Thus, to allow growth on methanol, we tested native or engineered enzymes originating from various organisms that were previously described to catalyze NAD-dependent methanol oxidation: *Bacillus methanolicus, Geobacillus staerothermophilus, Corynebacterium glutamicum,* and *Cupriavidus necator* (Wenk et al., 2020; Wu et al., 2016). Additionally, we tested two native alcohol dehydrogenases from *P. putida*: *adhP* and *pedE.* Through BLAST analyses, we discovered that the *adhP* gene encodes a homolog of the MDH of *C. glutamicum* (Identity: 41.5%). The *pedE* gene encodes a pyrroloquinoline quinone (PQQ) dependent alcohol dehydrogenase, that showed activity towards methanol as a substrate (Wehrmann et al., 2017). For the oxidation of formaldehyde to formate, we overexpressed the *fdhA* gene of *P. putida*, encoding a NAD-dependent formaldehyde dehydrogenase. The various MDH candidate genes and *fdhA* were cloned into pC1, creating pM. We transformed C1-S- Aux with the various pM plasmids to assess which MDH candidate can sustain the highest flux and therefore growth. All candidates were able to restore growth of the C1-S-Aux strain on glucose, glycine, and methanol, albeit with different growth patterns (Figure S1). The MDH from *C. necator* was able to sustain the best growth (10 hrs. doubling time) with the shortest lag phase (Figure 2b). Next, growth was fastest restored using the MDH from *B. methanolicus* and *C. glutamicum*, followed by *adhP* and *G. stearothermophilus.* The PQQ – dependent alcohol dehydrogenase from *P. putida* was able to sustain growth (11 hrs. doubling time) despite its long lag phase compared to the strains expressing a NAD-MDH. As far as we know, this is the first time a functional PQQ – dependent enzyme has been demonstrated for engineered methanol oxidation. Methanol oxidation using PQQ instead of NAD^+^ as an electron acceptor can be potentially advantageous as it has a larger thermodynamic driving force (Cotton et al., 2020). NAD-MDH activity is notorious for being a thermodynamic, as well as a kinetic bottleneck for synthetic methylotrophy (Woolston et al., 2018). Even though we showed proof of principle for the PQQ-dependent operation of an MDH, the best NAD-MDH results in faster growth in C1-S-Aux. Still, further optimization of PQQ-MDHs, including for example upregulation of native *P. putida* PQQ biosynthesis, can possibly sustain faster growth rates and may be beneficial to support full methylotrophy. However, in this work, we further proceeded with the best-performing NAD-MDH from *C. necator* to demonstrate the potential of the rGlyP for synthetic methylotrophy in *P. putida* (Figure 3b). Just as with formate, methanol became essential for the growth of C1-S-Aux and omitting it from the medium resulted in no growth. The observed growth (∼10 hrs. doubling time) was slower than growth of the control with formate (∼4 hrs.), which can be expected as methanol oxidation is rather slow. Moreover, overexpression of pM seems to lead to slower growth rates and a shorter exponential phase which is also observed in the controls containing pM and supplemented with serine. This could potentially be related to the metabolic burden of the plasmid containing a high number of genes. Labeling experiments with ^13^C-methanol further confirmed the combined activity of the M, C1, and C3 modules. As expected, serine and histidine were both labeled once (Figure 3c).

**Figure 3.**
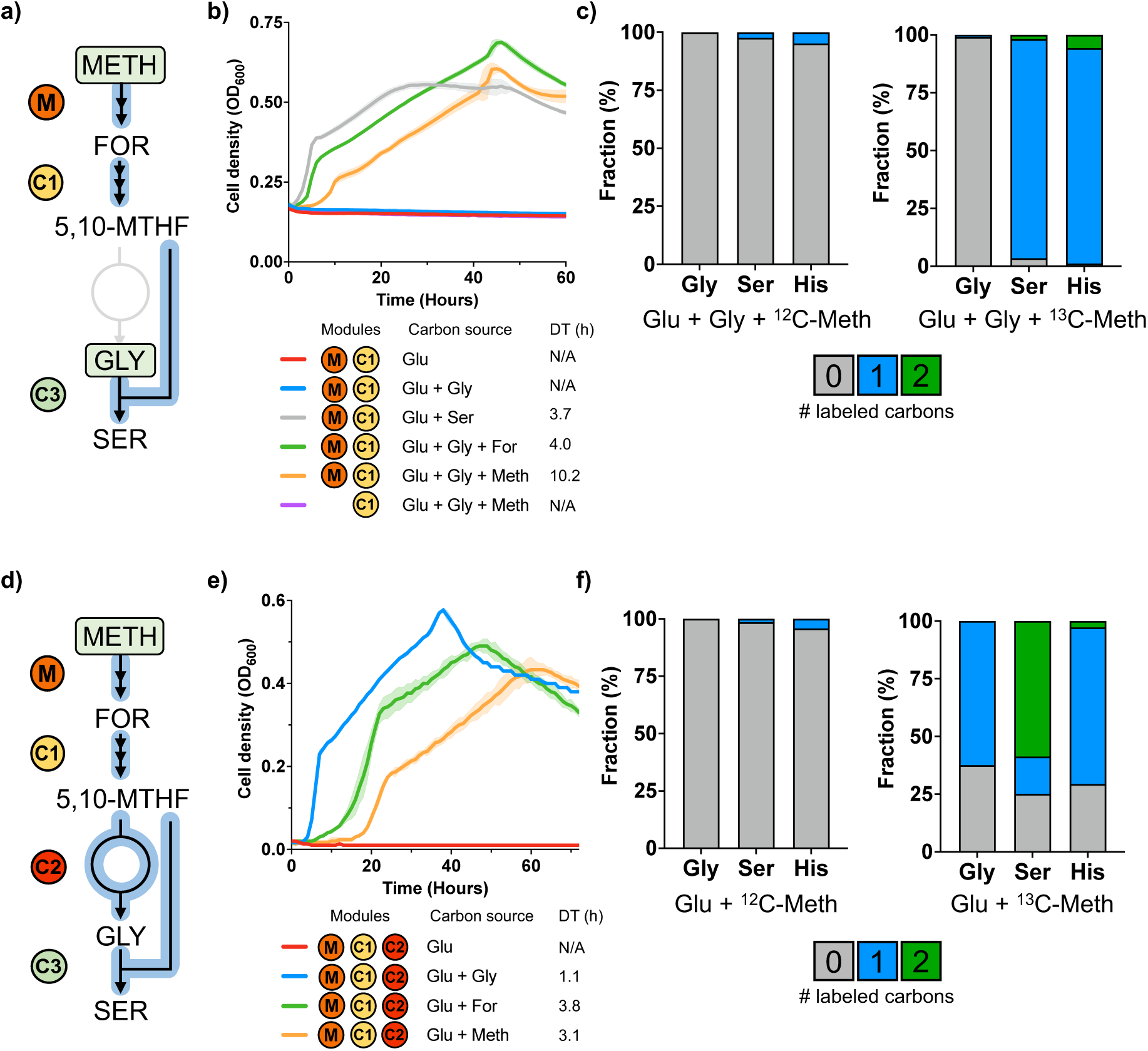
Methanol-dependent growth. **a)** Selection of the M and C1 modules in C1-S Aux. Growth is only possible when serine is supplemented to the medium or if both glycine and formate or methanol are present. **b)** Growth of C1-S Aux expressing the MDH CT4-1 from *Cupriavidus necator*. Growth is only possible upon overexpression of the M and C1 module, which replenishes the cellular C1-precursor molecules. **c)** ^13^C labeling experiments confirm that cellular C1-moieties are produced exclusively from methanol **d)** Selection of the combined effort of the M, C1, and C2 modules in C1-G-S Aux **e)** Growth of C1-G-S Aux. Growth can solely be restored when glycine or methanol or formate are added to the medium (at elevated CO_2_ levels through the addition of 100 mM sodium bicarbonate) **f)** ^13^C labeling experiments confirm that cellular C1-moieties, glycine, and serine are produced mostly from methanol and CO_2_. Abbreviations: (Gly) glycine, (Ser), serine, (His), histidine, (Meth), methanol, (For), formate (5,10-MTHF), 5,10-methylene-THF, (Glu), Glucose). Growth curves and labeling experiments are averages from biological triplicates.

We further tested methanol assimilation via the complete rGlyP till serine in the C1-G-S-Aux strain. Hereto, we tested pM with the engineered MDH from *C. necator* together with pC2. Through the combined effort of the M, C1, C2, and C3 modules, methanol together with CO_2_ could potentially provide the cell with the necessary glycine and serine. As expected, growth was restored upon expression of all the modules and the addition of the required C1-compounds (Figure 2e). We performed labeling experiments to confirm the activity of the combined modules (Figure 2f). We note that a small fraction of all glycine and serine is unlabeled. Glycine can be produced by the amination of glyoxylate through promiscuous aminotransferase enzymes. However, the isocitrate lyase (*aceA*) is deleted in C1-G-S Aux, making it unable to produce glyoxylate. We hypothesize that a latent unidentified reaction in *P. putida* can still produce glycine from threonine and was activated during growth of C1-G-S Aux on methanol, contributing to the unlabeled fraction (∼37% of glycine). Nonetheless, taking the growth and labeling patterns into account, we can still conclude that the methanol assimilation via the modules of the rGlyP carries the majority of flux in C1-G-S Aux.

### Extending the reductive glycine pathway with CO_2_ fixation

Apart from methanol oxidation, formate can be produced through CO_2_ reduction. FDHs usually convert formate to CO_2_, which is thermodynamically the most favorable direction. However, metal-dependent FDHs, using molybdenum or tungsten, can also serve in CO_2_ fixation pathways by reducing CO_2_ to formate (Cotton et al., 2018; Maia et al., 2017).

The genome of *P. putida* accounts for two native FDHs. The genes *fdoGHI - fdhE* (PP_0489 – 0492) encode a membrane-bound FDH, which possibly uses quinol as a redox cofactor. The second FDH (PP_2183-2186) is a soluble NAD-dependent molybdenum-containing FDH (Zobel et al., 2017). We reasoned that the molybdenum-containing NAD-FDH from *P. putida* could be able to reduce CO_2_ to formate and serve as an entry point for the rGlyP. To test this hypothesis, we constructed pCO_2_ containing PP_2183-2186 on a SEVA83b backbone.

We transformed C1-S-Aux with both pC1 and pCO_2_ and grew the strain in sealed bottles containing 20 mM glucose, 10 mM glycine, and CO_2_. The thermodynamics of CO_2_ reduction are highly unfavorable (Δ_r_G’^m^ = 22.5 kJ/mol, pH 7.5, ionic strength 0.25 M) (Flamholz et al., 2012). Therefore, we filled the headspace of the bottles with 50% (v/v) CO_2_ to push the reduction reaction. After a few weeks, we observed growth in one of the bottles. Reinoculation of this strain in fresh media with glucose, glycine, and 50% CO_2_ enabled immediate growth (Data not shown). We cultivated this strain, termed C1-S-Evo, in a range of different CO_2_ concentrations, from ambient (0.04%) to 50%, to examine the CO_2_ dependency of this strain. We observed that growth was highly dependent on the concentration of CO_2_ added to the headspace (Figure 4b). Fast growth was observed when the headspace was filled with 20 – 50% CO_2_, with doubling times ranging from 13.7 down to 6.0 hours, respectively. Growth still occurred at 10% CO_2_, albeit with a lower doubling time (39.7 hours). The clear dependency of the growth phenotype on the CO_2_ concentration is likely related to the thermodynamic driving force of CO_2_ reduction by FDH, which can be improved by increasing CO_2_ concentrations. Growth was not observed when the pCO_2_ plasmid was omitted, indicating that growth solely relies on the overexpression of FDH. Moreover, we noticed that this strain was able to grow at ambient CO_2_ levels (Figure 4b, gray line). We hypothesized that the respiration of glucose (still proceeding to supply other biomass components than C1) increases the CO_2_ concentration in the headspace, driving formate biosynthesis. To test this hypothesis, we cultivated the strain in a closed and open environment, wherein the latter the CO_2_ can freely escape. In both instances, growth occurred similarly (Figure S2). This indicates that the CO_2_ present in the air or derived from glucose respiration is enough to drive the reverse reaction towards formate production. To elucidate what changes had occurred during evolution to allow these CO_2_-fixing phenotypes, the pC1 and pCO_2_ plasmids of three growing isolates were sequenced.

**Figure 4.**
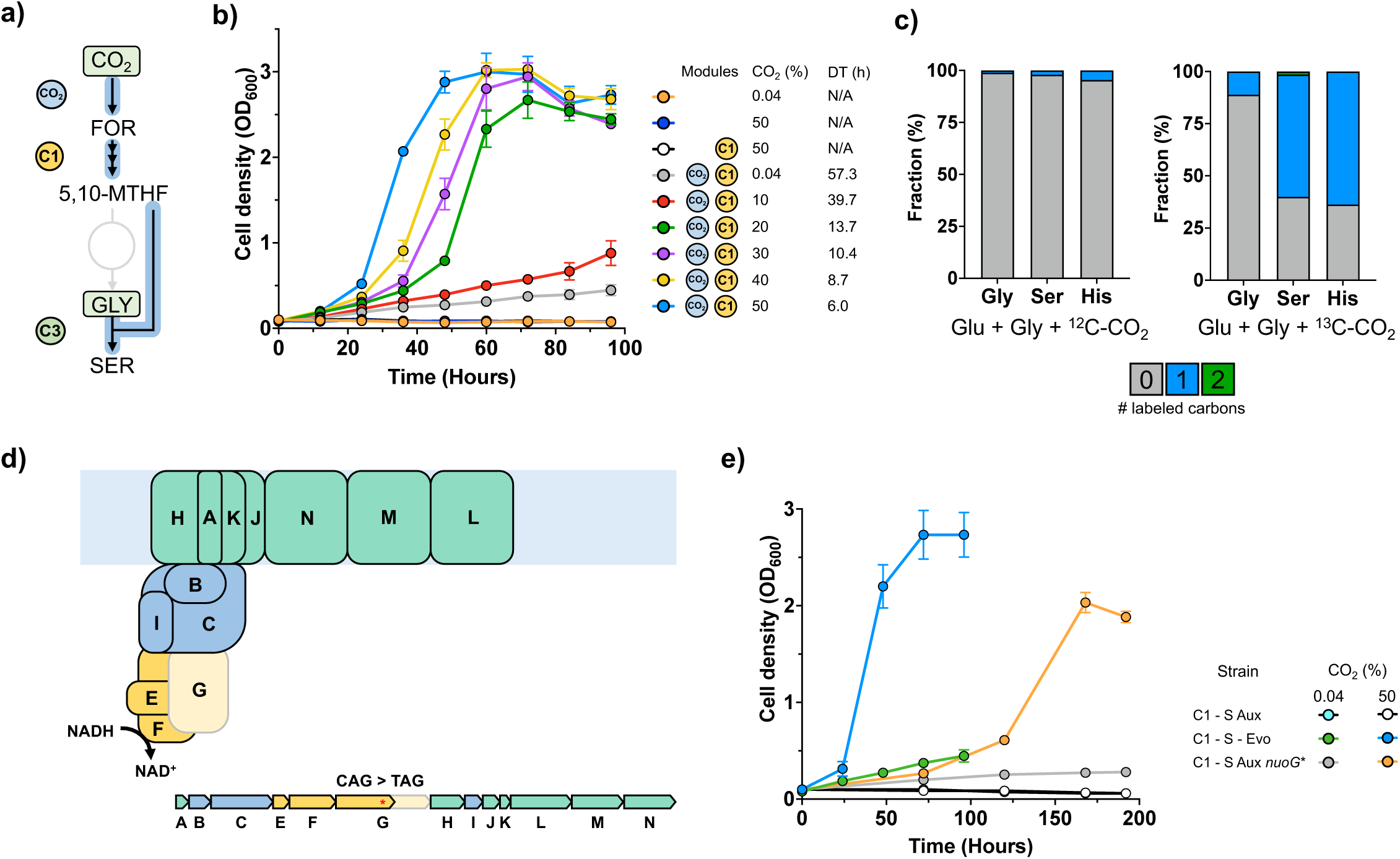
CO_2_-dependent growth. **a)** Selection of the CO_2_ and C1 modules in C1-S-Aux/Evo. Growth is only possible when serine is supplemented to the medium or if both glycine and CO_2_ are present. **b)** Growth of C1-S-Evo in different CO_2_ concentrations. Growth is only possible upon overexpression of the CO_2_ and C1 module, which replenishes the cellular C1-moieties. Growth is heavily dependent on the increasing CO_2_ concentrations. **c)** ^13^C labeling experiments confirm that cellular C1-moieties are produced mostly from CO_2_. **d)** Localization of the nuoG protein within complex I of the electron transport chain. The acquired stop codon (CAG>TAG) during evolution allows translation of only half the NuoG protein. Figure adapted from (Chadwick et al., 2018). **e)** Retro engineering of the TAG stop codon in the *nuoG* gene in C1-S-Aux. Growth was restored due to this single mutation at ambient (0.04) and 50% (v/v) CO_2_ by expressing the C1 and mutated CO_2_ module. Abbreviations: Abbreviations: (Gly) glycine, (Ser), serine, (His), histidine, (For), formate, (5,10-MTHF), 5,10-methylene-THF, (Glu), Glucose. Growth curves and labeling experiments are averages from biological triplicates.

We discovered a point mutation in the -35 box of the promoter of the CO_2_ module, lowering its expression by 31.5-fold based on GFP-fluorescence, indicating that the initial expression level of the promoter was too high (Figure S3). We transformed C1-S-Aux with pC1 and the mutated pCO_2_ plasmid, to analyze if this mutation was the sole cause for growth. However, no immediate growth was observed (Data not shown). Therefore, genomic alterations likely contributed to establish this CO_2_-dependent growth phenotype. We sequenced the genomes of three purified colonies from the evolved population and discovered four common mutations in all three isolates (Table S3). The most noticeable mutation was the introduction of a stop codon in the middle of the *nuoG* gene encoding the G subunit of the NADH-quinone oxidoreductase (complex I) in the electron transport chain. This complex is directly responsible for NADH oxidation and is needed to regenerate NAD^+^ for glucose respiration. The occurred mutation allows translation of only half of *nuoG*, probably either interrupting or decreasing the NADH oxidation activity complex I (Figure 4d). FDH needs high levels of NADH and CO_2_, to push the thermodynamically challenging reaction towards formate formation. Usually, *P. putida* and other organisms maintain a relatively low NADH/NAD^+^ ratio, which likely prevents CO_2_ reduction by FDH in the unevolved strain. Thus, we hypothesize that this mutation causes a redox perturbance that increases NADH levels for CO_2_ reduction., We reverse-engineered the stop codon mutation in *nuoG* in C1-S-Aux to test the influence of this mutation and transformed the strain with pC1 and the mutated pCO_2_ plasmid. Surprisingly, with only this mutation, growth could occur at ambient and 50% CO_2_ without the need for evolution, showcasing the beneficial effect of this mutation (Figure 4e). However, growth was still lagging compared to the evolved strain, so some of the other mutations may have contributed partly to the phenotype.

To further prove CO_2_ reduction and its entry into the rGlyP, we performed labeling experiments with ^13^C–CO_2_. Here, serine was expected to be labeled once completely but was labeled for only ∼60% (50% minus the labeled glycine). We demonstrated earlier that CO_2_ released from glucose respiration is enough to drive the reverse FDH reaction resulting in slow growth. Therefore, it is likely that a mixture of labeled, as well as unlabeled intracellular CO_2_ originating from glucose, is being fixed by the FDH, leading to mixed labeling patterns. This pattern is repeated for histidine, which should theoretically be fully labeled once. To estimate the intracellular labeling status of CO_2_, we analyzed the proteinogenic amino acids proline and arginine (Figure S4). In the biosynthesis of arginine, one CO_2_ is added through carboxylation, which is not present in proline. All other carbons of proline and arginine have the same origin (Gleizer et al., 2019) Therefore, the difference in labeling between these two amino acids can be used to estimate the labeling of intracellular CO_2_. This analysis showed that ∼76 % of intracellular CO_2_ is labeled. So, one would expect somewhat higher labeling than 60% for serine. This may indicate some uncertainty in the method to determine intracellular CO_2_ or some (small) contribution of other latent pathways to C1-biosynthesis other than CO_2_ reduction. However, considering the dependency of growth on CO_2_ and labeling patterns (∼50- 60% for serine and histidine), we can conclude that CO_2_ fixation via the reverse FDH reaction is the main contributor to C1-biosynthesis.

After establishing CO_2_-dependent growth in the C1-S-Evo strain, we aimed to achieve the same in C1-G-S-Aux. For this purpose, we refactored the C1-S-Evo strain, instead of introducing the necessary mutations in C1-G-S-Aux. We deleted the *ltaE* and *aceA* genes to prevent glycine formation and reintroduced the C2 module in the genome. This strain, termed C1-G-S-Evo, was transformed with pC1 and the mutated pCO_2_, and growth was assessed on 20 mM glucose and 50% CO_2_. The strain expressing all three modules was able to sustain growth, albeit very slowly and with a low biomass yield (Figure S5). Although promising, further optimizations are needed to build towards strains that can generate all biomass from CO_2_ via FDH.

## DISCUSSION

In this study, we successfully laid the foundation for synthetic C1 – metabolism in the industrial workhorse *P. putida*. We were able to demonstrate formate, methanol, and CO_2_ assimilation through heterologous expression of the core modules of the rGlyP in growth-coupled, auxotrophic selection strains. We show functional expression of all key modules of the rGlyP until serine using both formate and methanol as substrate. The demonstrated conversion in C1- G-S-Aux of methanol and formate into glycine and serine (together forming ∼11% of cellular biomass (Claassens et al., 2019)) provides a strong basis for full formatotrophy and methylotrophy in *P. putida*. Full C1-dependent growth on both substrates can likely be achieved by a combination of genomic integration and further fine-tuning of the enzymes of the independent modules. Then, a combination of rational engineering and adaptive laboratory evolution can be used to optimize the complete metabolic network. The establishment of full C1-dependent growth in *P. putida* will open many avenues for C1-based industrial biotechnology, given the many attractive properties, and demonstrated production pathways available for this bacterium.

Establishing full formatotrophy in *P. putida* can benefit from its native catalytically fast, metal- dependent FDH, to provide energy to run the rGlyP. This enzyme is natively lacking in *E. coli,* and heterologous expression of metal-dependent FDHs in *E. coli* has been proven challenging. So far, full formatotrophic growth in *E. coli* was only demonstrated by heterologously expressing a slower non-metal-dependent FDH, likely limiting formatotrophic growth rates (Bar-Even et al., 2013; Gleizer et al., 2019; Kim et al., 2020).

The first demonstration of synthetic methylotrophy in *P. putida,* shown in this work, also showcases a specific promising feature of *P. putida* to engineer methanol conversion, which could proceed through PQQ-dependent MDH activity. Methanol oxidation is frequently pinpointed as the major bottleneck during the ongoing efforts to establish synthetic methylotrophy in other organisms (Antoniewicz, 2019; Wang et al., 2020). The NAD- dependent enzymes can lead to higher biomass yields but come with disadvantageous thermodynamics. The PQQ-dependent enzymes do cause a slight reduction in biomass yield, but their higher thermodynamics could lead to higher growth rates (Claassens et al., 2022; Cotton et al., 2020; Whitaker et al., 2015). So far, none of all the published metabolic engineering efforts towards synthetic methylotrophy have demonstrated the engineered expression of a PQQ-dependent MDH. As we demonstrate here, *P. putida* is a promising host to establish PQQ-dependent synthetic methylotrophy, due to its native PQQ biosynthesis and PQQ-alcohol dehydrogenase that could execute PQQ-MDH activity. Although methanol- dependent growth in C1-S-Aux via PQQ-MDH is slower than via the NADH-MDH, these results establish the basis to further develop this industrially attractive phenotype. The native PQQ-MDH candidate enzyme used in this study could be further engineered through directed evolution toward higher specificity and activity on methanol. In summary, both NAD- and PQQ-dependent MDHs could be further explored to realize efficient and fast synthetic methylotrophy in *P. putida*.

This study shows for the first time engineered CO_2_ fixation via the reverse activity of FDH. In recent years, CO_2_ to formate reduction has been proposed for sustainable biotechnology as a promising feature for both engineered *in vitro* and *in vivo* CO_2_ fixation. However, so far experimental evidence of engineered CO_2_ fixation via FDH has only been shown for *in vitro* CO_2_ reduction. These *in vitro* studies were based on metal-dependent FDH enzymes, from for example *C. necator* and *Rhodobacter capsulatus* (Hartmann & Leimkühler, 2013; X. Yu et al., 2017). The high *in vitro* catalytic rates found for CO_2_ reduction to formate of these metal- dependent FDH enzymes (as opposed to slower non-metal-dependent FDHs) are a promising indication that *in vivo* synthetic CO_2_ fixation via metal-dependent FDH is achievable. However, engineered *in vivo* activity from CO_2_ to formate by a metal-dependent FDH was not shown yet, possibly due to the complexity of overexpressing metal-dependent FDHs, which typically require chaperones and consist of multiple subunits. In addition, the reduction of CO_2_ to formate by FDH is thermodynamically relatively challenging, likely requiring a very high substrate (CO_2_ and NADH) to product (formate and NAD^+^) ratio within the cell. We overcome these limitations by expressing the native metal-dependent FDH at elevated CO_2_ levels, enabling for the first time FDH-mediated CO_2_ reduction activity *in vivo.* By using a growth-coupled selection strategy, we show that the reaction can carry sufficient flux to supply the cell with the C1-precursors and the beta-carbon of serine (forming ∼4% of cellular biomass) (Claassens et al., 2019). However, this growth phenotype required short-term laboratory evolution, which resulted in fine-tuning of FDH expression and an essential early stop codon mutation in the NuoG subunit of complex I in the electron transport chain. This likely increased the NADH/NAD^+^ ratio, facilitating the thermodynamics of CO_2_ reduction to formate. In *E. coli*, it has been shown that a similar deletion rendered complex I non-functional, most likely resulting in an unbalanced NADH/NAD+ ratio (Falk-Krzesinski & Wolfe, 1998). This indicates that the establishment of CO_2_ fixation via FDH in *P. putida* and potentially other organisms require modulation of the NADH/NAD^+^ ratio. Alternatively, metal-dependent NADPH-FDHs could be further developed for CO_2_ reduction, as cells typically maintain a higher NADPH/NADP^+^ ratio compared to NADH/NAD^+^ (Calzadiaz-Ramirez & Meyer, 2022). Demonstrating efficient CO_2_ reduction until serine in the C1-G-S-auxotroph was still not achieved. This likely reflects the more challenging redox requirements to realize this higher flux towards all C1-precursors, glycine, and serine in the cell. The reduction of CO_2_ to formate could probably be further improved by further redox-factor engineering approaches mentioned above and possibly with engineered or heterologous metal-dependent FDH candidates, which could be potentially even faster.

Overall, achieving growth with CO_2_ as the sole carbon source (synthetic autotrophy) in *P. putida* via the rGlyP would be a very promising feature for sustainable industrial biotechnology. This would also require a suitable inorganic electron donor, for which hydrogen is an interesting candidate as it can be produced very efficiently from renewable electricity (Claassens et al., 2018). The soluble oxygen tolerant NAD-reducing hydrogenase from *C. necator* has already been successfully expressed *in vivo* in *P. putida* as an NADH regeneration system for product synthesis (Lonsdale et al., 2015). This NADH regeneration system combined with the here established CO_2_ reduction could enable synthetic autotrophy in *P. putida* based on the rGlyP. If the FDH activity could be increased, which based on *in vitro* data could be faster than Rubisco (Cotton et al., 2018), the rGlyP could possibly be a kinetically faster alternative to the naturally, ubiquitous Calvin cycle. This, together with the lower ATP costs of the rGlyP versus the Calvin Cycle makes it an attractive pathway to explore for synthetic autotrophy in *P. putida* and other organisms.

Overall, this work widens the possibilities for engineered C1–assimilation based on the versatile rGlyP and shows the suitability of *P. putida* for C1-based biotechnology through lifestyle engineering. The successful establishment of synthetic C1-assimilation in *P. putida* is a key step to realize a truly sustainable C1-based biotechnology.

## METHODS

### Plasmids, primers, and strains

All strains and plasmids used in the present study are listed in Table S1. Primers used for plasmid construction and gene deletions are listed in Table S2.

### Bacterial strains and growth conditions

*P. putida* and *E. coli* cultures were incubated at 30°C and 37°C respectively. For cloning purposes, both strains were propagated in Lysogeny Broth (LB) medium containing 10 g/L NaCl, 10 g/L tryptone, and 5 g/L yeast extract. For the preparation of solid media, 1.5% (w/v) agar was added. Antibiotics, when required, were used at the following concentrations: kanamycin (Km) 50 μg/ml, gentamycin (Gm) 10 μg/ml, chloramphenicol (Cm) 50 μg/ml and apramycin (Apra) 50 μg/ml. All growth experiments were performed using M9 minimal medium (Hartmans et al., 1989). Before the growth experiments, cells were pre-grown in non-selective M9 media, containing 10 mM glucose and 2 mM serine, before being transferred to selective growth conditions. In all growth experiments, precultured strains were washed twice in M9 media without a carbon source before transfer and inoculated at an OD_600_ of 0.1. Selective growth conditions consisted of unless otherwise indicated, 10 mM glucose + 5 mM glycine (C1-S-Aux) or 10 mM glucose (C1-G-S Aux) supplemented with relevant C1- substrates (30 mM formate or 500 mM methanol or 50% (v/v) CO_2_). For the formate and methanol experiments with C1-G-S-Aux, 100 mM sodium bicarbonate was added to the medium to push the GCS in the reverse direction. Plate reader experiments were carried out in 200 μL of M9 medium using a Synergy plate reader (Biotek). Growth (OD_600_) was measured over time using continuous linear shaking (567 cpm, 3mm) and measurements were taken every five minutes. For the CO_2_-dependent growth experiments, 100 mL glass bottles were filled for 10% (10 mL) with liquid M9 media. Subsequently, the headspace (90%) was filled with the desired CO_2_ concentration before autoclavation. Carbon sources and antibiotics were added after sterilization and cultures were incubated in a rotary shaker at 200 rpm at 30°C. All growth experiments were performed in biological triplicates and the represented growth curves show the average of these triplicates.

### Plasmid construction

Plasmids were constructed using the standard protocols of the previously described SevaBrick Assembly (Damalas et al., 2020). All DNA fragments were amplified using Q5^®^ Hot Start High-Fidelity DNA Polymerase (New England Biolabs). To construct the C1 module, DNA fragments of the *fhs*, *fchA,* and *folD* genes from *Clostridium ljunghdahlli* DSM13528 were codon-optimized with the JCat tool (Grote et al., 2005) and synthesized through Genescript. The MDH genes from *Geobacillus. staerothermophilus, Bacillus. methanolicus, Corynebacterium glutamicum,* and an engineered variant from *Cupriavidus. Necator* (Wu et al., 2016) were codon-optimized and synthesized by IDT (Integrated DNA Technologies) (TableS4). All genes in this study were expressed under the control of a strong constitutive promoter (BBa_J23100) and RBS (BBa_B0034). All plasmids were transformed using heat shock in chemically competent *E. coli* DH5α λpir and selected on LB agar with corresponding antibiotics. Colonies were screened through colony PCR with Phire Hot Start II DNA Polymerase (Thermo Fisher Scientific). Isolated plasmids were verified using Sanger sequencing (MACROGEN inc.) and subsequently transformed into *P. putida* via electroporation.

### Genome modification

Genomic deletions in this study were performed using the protocol previously described by Wirth et al. (2020). Homology regions of ± 500 bp were amplified up and downstream of the target gene from the genome of *P. putida* KT2440. Both regions were cloned into the non-replicative pGNW vector and propagated in *E. coli* DH5α λpir. Correct plasmids were transformed into *P. putida* by electroporation and selected on LB + Km plates. Successful co-integrations were verified by PCR. Hereafter, co-integrated strains were transformed with the pQURE6-H, and transformants were plated on LB + Gm containing 2 mM 3- methylbenzoic acid (3-mBz). This compound induces the XylS – dependent Pm promoter, regulating the *I- Sce*I homing nuclease that cuts the integrated pGNW vector. Successful gene deletions were verified by PCR and Sanger sequencing (MACROGEN inc). Hereafter, the pQURE6-H was cured by removing the selective pressure and its loss was verified by sensitivity to gentamycin.

### Promoter characterization

*P. putida* strains expressing GFP under the normal or mutated J23100 promoter or containing an empty vector were grown in biological triplicates in M9 medium + 10 mM glucose. Cell density (OD600) and GFP fluorescence (excitation 485 nm, emission 512 nm, gain 50) were measured using a Synergy plate reader (Biotek) over time using continuous linear shaking, and measurements were taken every five minutes. The promoter strength was quantified based on fluorescence normalized per cell density (RFU/OD_600_) after 20 hours of cultivation when cells had reached the stationary phase. Values were corrected for background fluorescence of cells without GFP.

### Whole-genome sequencing

Genomic DNA of the evolved mutants was isolated from LB overnight cultures using the GenElute™ Bacterial Genomic DNA Kit (Sigma-Aldrich St. Louis, MO). The extracted DNA was evaluated by gel electrophoresis and quantified by a NanoDrop spectrophotometer (Thermo Fisher Scientific). Samples were sent for Illumina sequencing to Novogene Co. Ltd. (Beijing, China). Raw Illumina reads were trimmed for low quality and adapters with fastp (v0.20.0). Mutations were identified by comparing the reads to the annotated reference genome of *Pseudomonas putida* KT2440 (GCF_000007565.2) using breseq (v0.35.5) (Barrick et al., 2014).

### Carbon labeling

For stationary isotope tracing of the proteinogenic amino acids, cultures were grown in 10 ml of M9 media under the previously described experimental conditions. Media was composed of unlabeled (glucose and glycine) and labeled (formate – ^13^C, methanol – ^13^C, and ^13^CO_2_) carbon sources. Cells were cultivated in two successive cultures with labeled or unlabeled carbon to diminish the labeling effects of the preculture. Hereafter, the approximate cell volume was harvested that has the cellular biomass roughly equivalent to 1 mL with an OD600 of 1 was taken and pelleted down. The pellet was washed with 1 mL of pure water and pelleted down again. The pellet was resuspended in 1 ml of 6N HCL and incubated for 24 hours at 95 °C. Then, the caps were opened, allowing evaporation under continuous airflow. The resulting pellet was resuspended in 1 mL of pure water and centrifugated for 5 minutes at full speed to remove residual particles. The hydrolysate was analyzed using ultra-performance liquid chromatography (UPLC) (Acquity, Waters) using an HSS T3 C18-reversed-phase column (Waters). The mobile phases were 0.1% formic acid in H2O (A) and 0.1% formic acid in acetonitrile (B). The flow rate was 400 μL/min and the following gradient was used: 0–1 min 99% A; 1–5 min gradient from 99% A to 82%; 5–6 min gradient from 82% A to 1% A; 6–8 min 1% A; 8–8.5 min gradient to 99% A; 8.5–11 min – re-equilibrate with 99% A. Mass spectra were acquired using an Exactive mass spectrometer (MS) (Thermo Scientific) in positive ionization mode. Data analysis was performed using Xcalibur (Thermo Scientific). The identification of amino acids was based on retention times and m/z values obtained from amino acid standards (Sigma-Aldrich)

## Acknowledgments

We are grateful to Iame Alves Guedes and Sara Cantera Ruiz de Pellon for their invaluable help with the CO_2_-dependent experiments. We thank Bart Nijse for the analysis of the whole genome sequencing. This work was financed by the European Union’s Horizon2020 Research and Innovation Program under grant agreement Nos. 635536 (EmPowerPutida) and 730976 (IBISBA) to V.A.P.M.d.S.

## Author contributions

L.B. conceived the study, performed growth experiments, and metabolome sampling; L.B., S.W, and N.J.C designed the experiments; S.W. analyzed the ^13^C tracer analysis; L.B. wrote the initial draft; L.B., S.W., N.J.C and V.A.P.M.d.S. reviewed and edited the manuscript; N.J.C. and V.A.P.M.d.S. provided supervision. V.A.P.M.d.S. arranged the funding.

## Correspondence

Correspondence and request for materials should be addressed to L.B., N.J.C, or V.A.P.M.d.S

## SUPPLEMENTARY FIGURES

**Figures S1.**
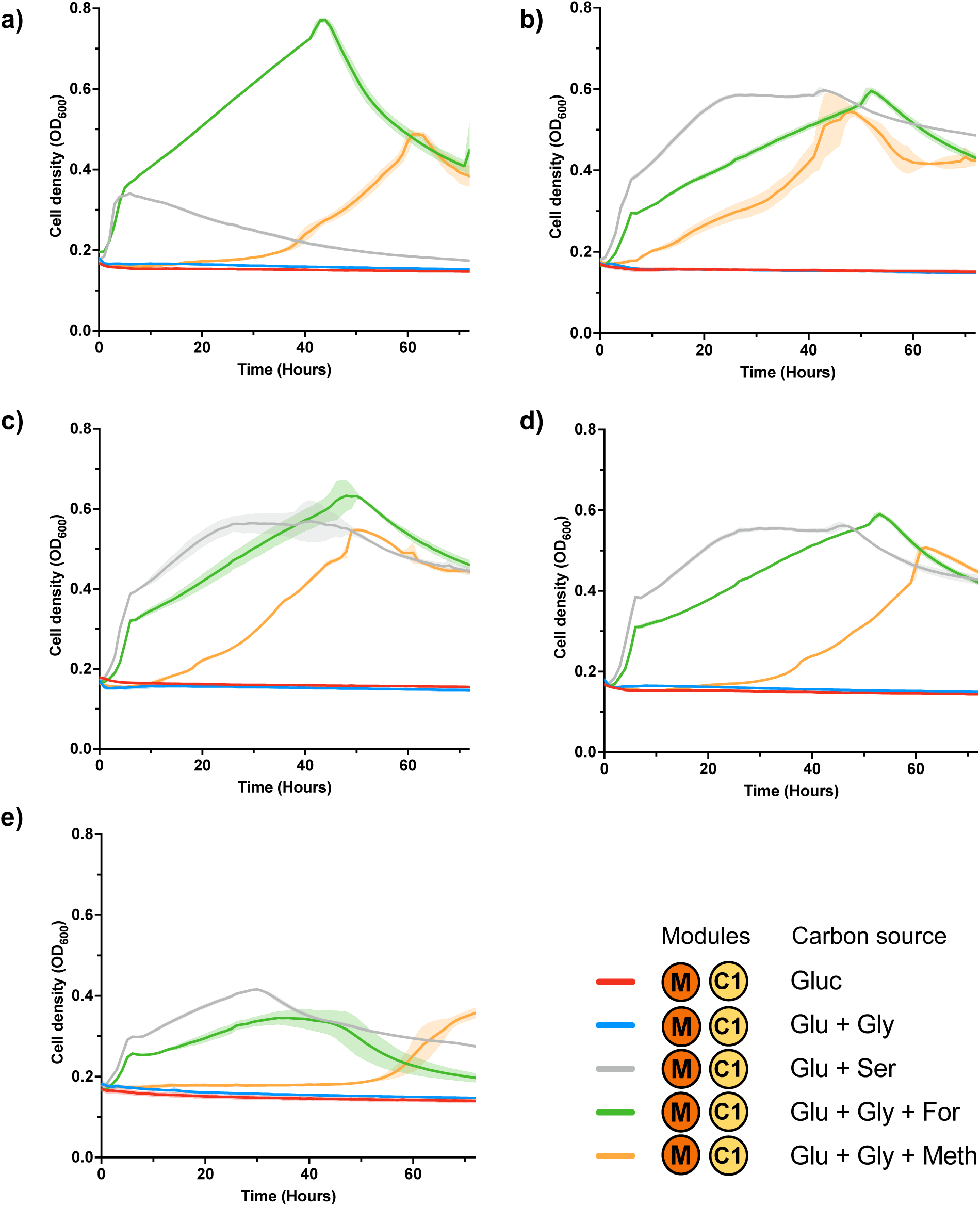
Methanol-dependent growth in C1-S-Aux with different methanol dehydrogenases. **a)** *G.stearothermophilus*, **b)** *B. methanolicus,* **c)** *C.glutamicum*, **d)** *adhP P. putida*, **e)** *pedE P. putida.* Growth curves are averages from biological triplicates.

**Figure S2.**
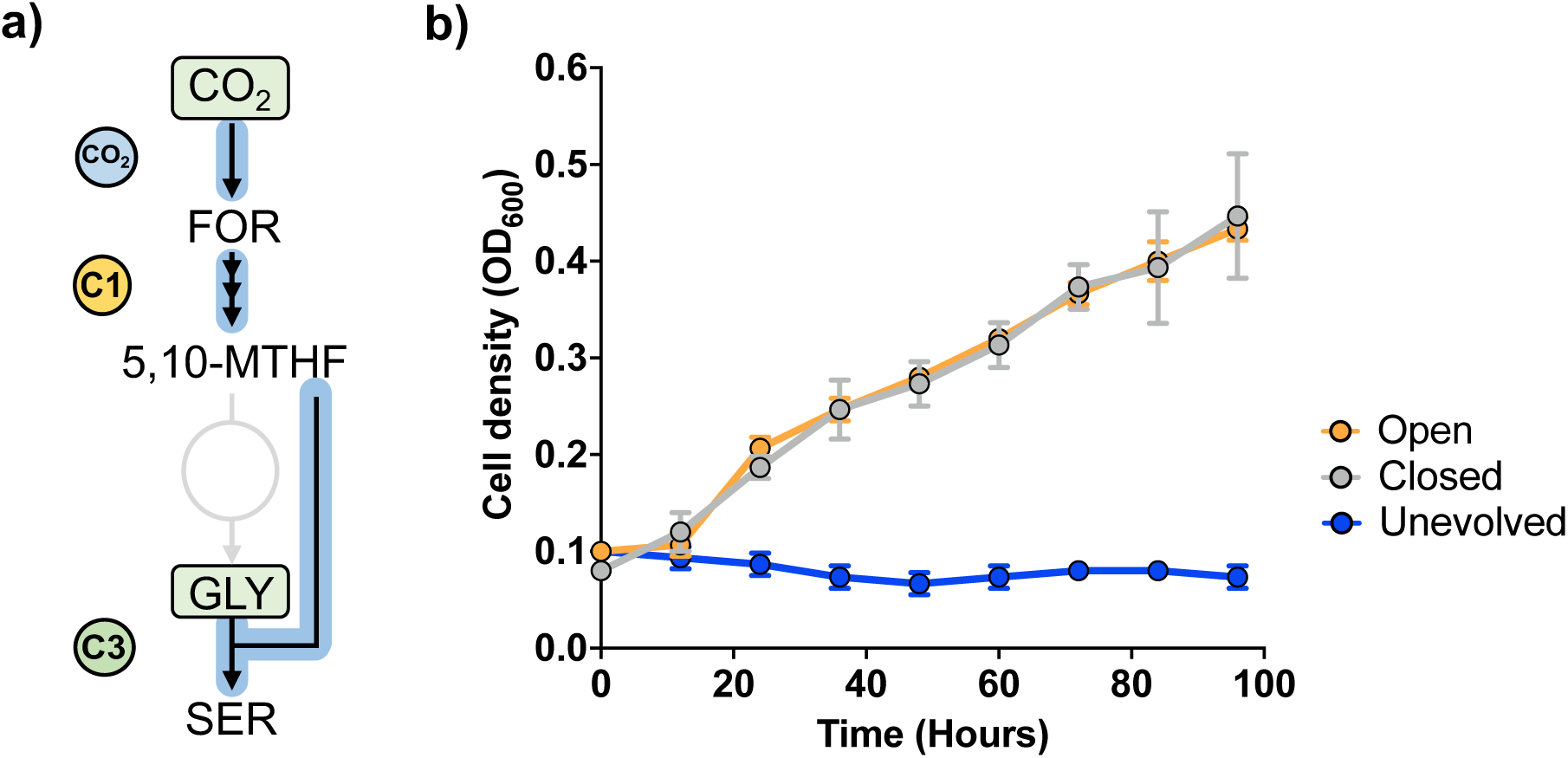
CO_2_ - dependent growth at ambient CO_2_ with different cultivation systems. **a)** Selection of the CO_2_-dependent growth in C1- S-Evo. b) Growth patterns of C1-S-Evo cultivated in closed anaerobic bottles and open falcon tubes where the CO_2_ can freely escape. The unevolved C1-S Aux strain was taken as a control. Growth curves are averages from biological triplicates.

**Figure S3.**
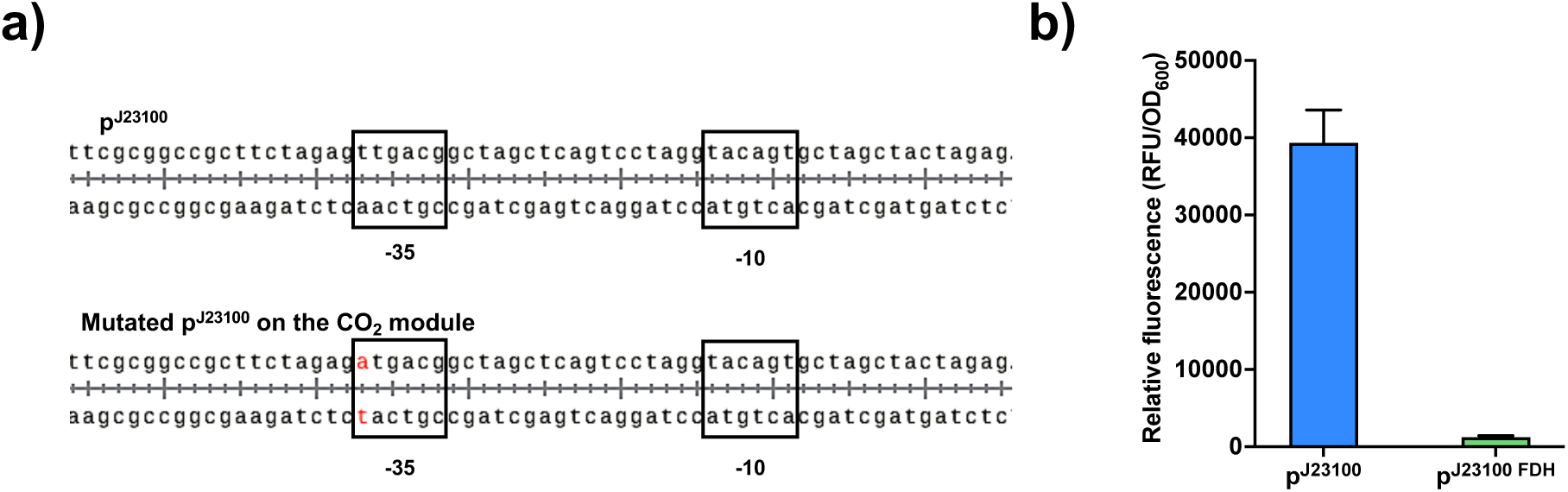
Characterization of the mutated CO_2_ module during short-term evolution. a). Promoter sequences of the J23100 promoter and the mutated CO_2_ module. b). Characterization of the mutated J23100 promoter of the CO_2_ module using fluorescence as output. Relative fluorescence averages were taken from biological triplicates.

**Figure S4.**
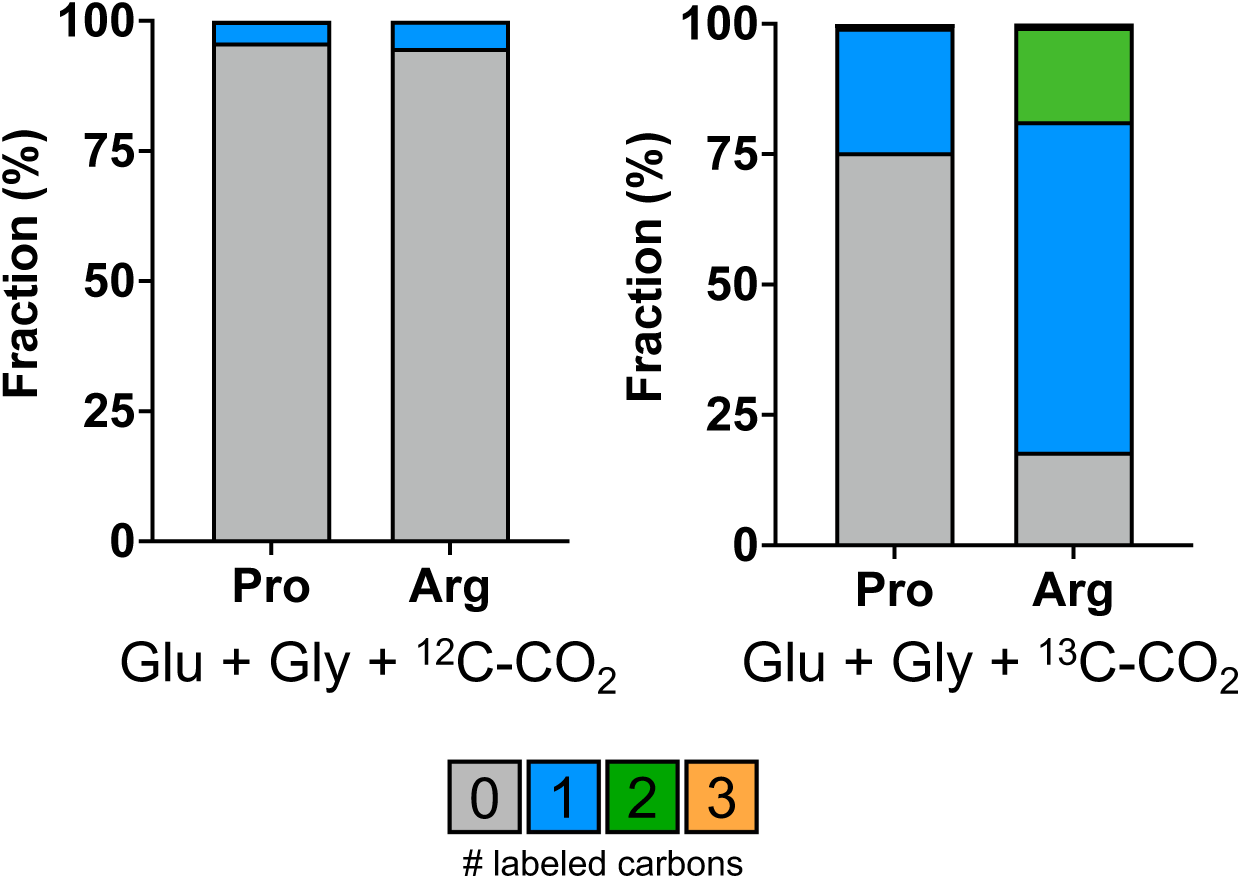
^13^C labeling of proline (Pro) and arginine (Arg) to determine intracellular CO_2_ levels in C1-S-Evo. Labeling experiments are averages from biological triplicates.

**Figure S5.**
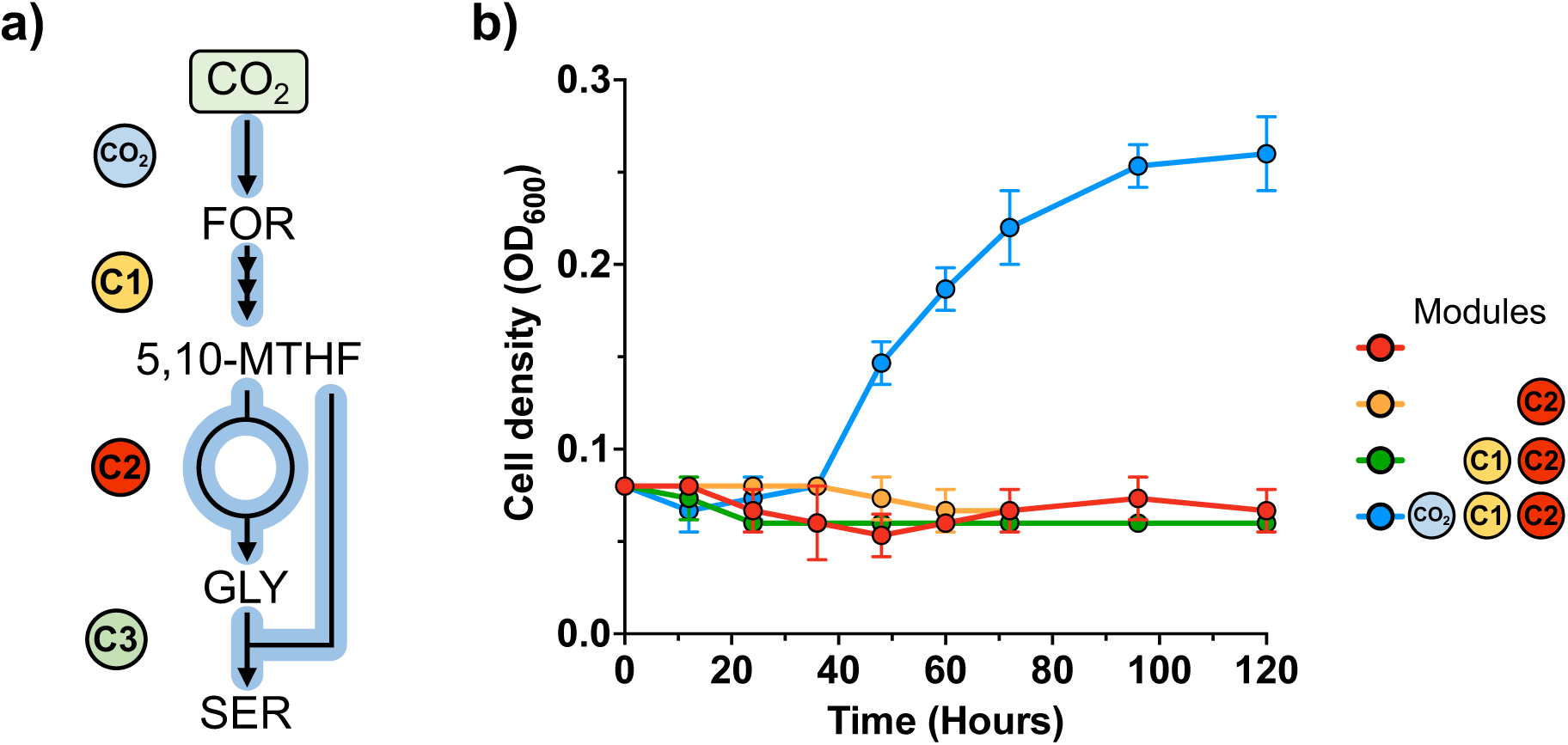
CO_2_ - dependent growth in C1-G-S-Evo. **a)** Selection of the combined effort of the CO_2_, C1, and C2 modules in C1-G-S- Evo. **b)** Growth was solely restored when the CO_2_, C1, and C2 modules are expressed. Growth curves are averages from biological triplicates.

## SUPPLEMENTARY TABLES

**Table S 1.** Strains and plasmids used in the present study.

| Strains | Description | Source |
| --- | --- | --- |
| ***Escherichia coli*** |  |  |
| Dh5α | Cloning host: F-λ-endA1 glnX44(AS) thiE1 recA1 relA1 spoT1 gyrA96(NalR) rfbC1 deoR nupG Φ80(lacZΔM15) Δ(argF-lac)U169 hsdR17(rK mK ) | (Grant et al., 1990) |
| CC118λpir | Cloning host: araD139 Δ(ara-leu)7697 ΔlacX74 galE galK phoA20 thi<sup>-</sup> 1 rpsE rpoB(Rif<sup>R</sup>) argE(Am) recA1, λpir lysogen | (Herrero et al., 1990) |
| ***Pseudomonas putida*** |  |  |
| KT2440 | Wild-type strain; mt-2 derivative cured of the TOL plasmid pWW0 | (Bagdasarian et al., 1981) |
| C1-S-Aux | KT2440 Δ*gcvTPH-I* Δ*gcvTPH-II* Δ*serA* Δ*PP* 2533 | This work |
| C1-G-S Aux | KT2440 Δ*serA* Δ*PP* 2533 Δ*ltaE* Δ*aceA* | This work |
| C1-S— Evo | Evolved C1-S-Aux | This work |
| C1-G-S – Evo | C1-S-Aux – Evo Δ*ltaE* Δ*aceA* :*gcvTPH-I* | This work |
| C1-S-Aux: nuoG* | C1-S-Aux with the reintroduced TAG stop codon in *nuoG* | This work |
| **Plasmids** |  |  |
| pGNW | Plasmid for genome editing in Gram-negative bacteria: ori (R6K) neo, Km<sup>R</sup> | (Wirth et al., 2020) |
| pGNW – *gcvTPH-I* | pGNW for scarless deletion of *gcvTPH-I* | This study |
| pGNW – *gcvTPH-II* | pGNW for scarless deletion of *gcvTPH-II* | This study |
| pGNW – *serA* | pGNW for scarless deletion of *serA* | This study |
| pGNW – PP 2533 | pGNW for scarless deletion of PP 2533 | This study |
| pGNW – *ltaE* | pGNW for scarless deletion of *ltaE* | This study |
| pGNW – *aceA* | pGNW for scarless deletion of *aceA* | This study |
| pGNW – C2 module | pGNW for scarless insertion of *gcvTPH-I* | This study |
| pGNW – *nuoG* - TAG | pGNW to introduce a TAG stop codon in *nuoG* | This study |
| pQURE6-H | Helper plasmid; oriV(RK2), xylS, Pml-SceI; Gm<sup>R</sup> | (Volke et al., 2020) |
| pC1 | pSEVAb24 containing the *fhs*, *fchA* and *folD* genes from *Clostridium ljungdahlii* DSM 13528 :ori (pRO1600/ColE1), Km<sup>R</sup> | This study |
| pC2 | pSEVAb65 containing the *gcvT-I*, *gcvP-I* and *gcvH-I* genes of *Pseudomonas putida* : ori (RSF1010) neo, Gm<sup>R</sup> | This study |
| pM- *G.stae* | pSEVAb24 containing the *mdh* from *Geobacillus stearothermophilus*, *fhdA* from *Pseudomonas putida* and *fhs*, *fchA* and *folD* genes from *Clostridium ljungdahlii* DSM 13528 :ori (pRO1600/ColE1), Km<sup>R</sup> | This study |
| pM-*B.met* | pSEVAb24 containing *mdh3* from *Bacillus methanolicus*, *fhdA* from *Pseudomonas putida* and *fhs*, *fchA* and *folD* genes from *Clostridium ljungdahlii* DSM 13528 :ori (pRO1600/ColE1), Km<sup>R</sup> | This study |
| pM-*C.nec* | pSEVAb24 containing *mdh CT4-1* from *Cupriavidus necator*, *fhdA* from *Pseudomonas putida* and *fhs*, *fchA* and *folD* genes from *Clostridium ljungdahlii* DSM 13528 :ori (pRO1600/ColE1) , Km<sup>R</sup> | This study |
| pM-*C.glu* | pSEVAb24 containing *adhP* from *Corynebacterium glutamicum*, *fhdA* from *Pseudomonas putida* and *fhs*, *fchA* and *folD* genes from *Clostridium ljungdahlii* DSM 13528 :ori (pRO1600/ColE1), Km<sup>R</sup> | This study |
| pM-*P.put* | pSEVAb24 containing *adhP* from *Pseudomonas putida*, *fhdA* from *Pseudomonas putida* and *fhs*, *fchA* and *folD* genes from *Clostridium ljungdahlii* DSM 13528 :ori (pRO1600/ColE1), Km<sup>R</sup> | This study |
| pM-*pedE* | pSEVAb24 containing *pedE* from *Pseudomonas putida*, *fhdA* from *Pseudomonas putida* and *fhs*, *fchA* and *folD* genes from *Clostridium ljungdahlii* DSM 13528 :ori (pRO1600/ColE1), Km<sup>R</sup> | This study |
| pCO<sub>2</sub> | pSEVAb83 containing PP_2183 – PP_2186 from *Pseudomonas putida* :ori (pBBR1), Apr<sup>R</sup> | This study |
| pCO<sub>2</sub>-mut | pSEVAb83 containing PP_2183 – PP_2186 from *Pseudomonas putida* with a mutated J23100 promotor :ori (pBBR1), Apr<sup>R</sup> | This study |
| pGFP | pSEVAb83 containing a GFP with a mutated J23100 promotor :ori (pBBR1), Apr<sup>R</sup> | This study |
| p-GFP-mut | pSEVAb83 containing a GFP with the J23100 promotor :ori (pBBR1), Apr<sup>R</sup> | This study |

**Table S 2.**
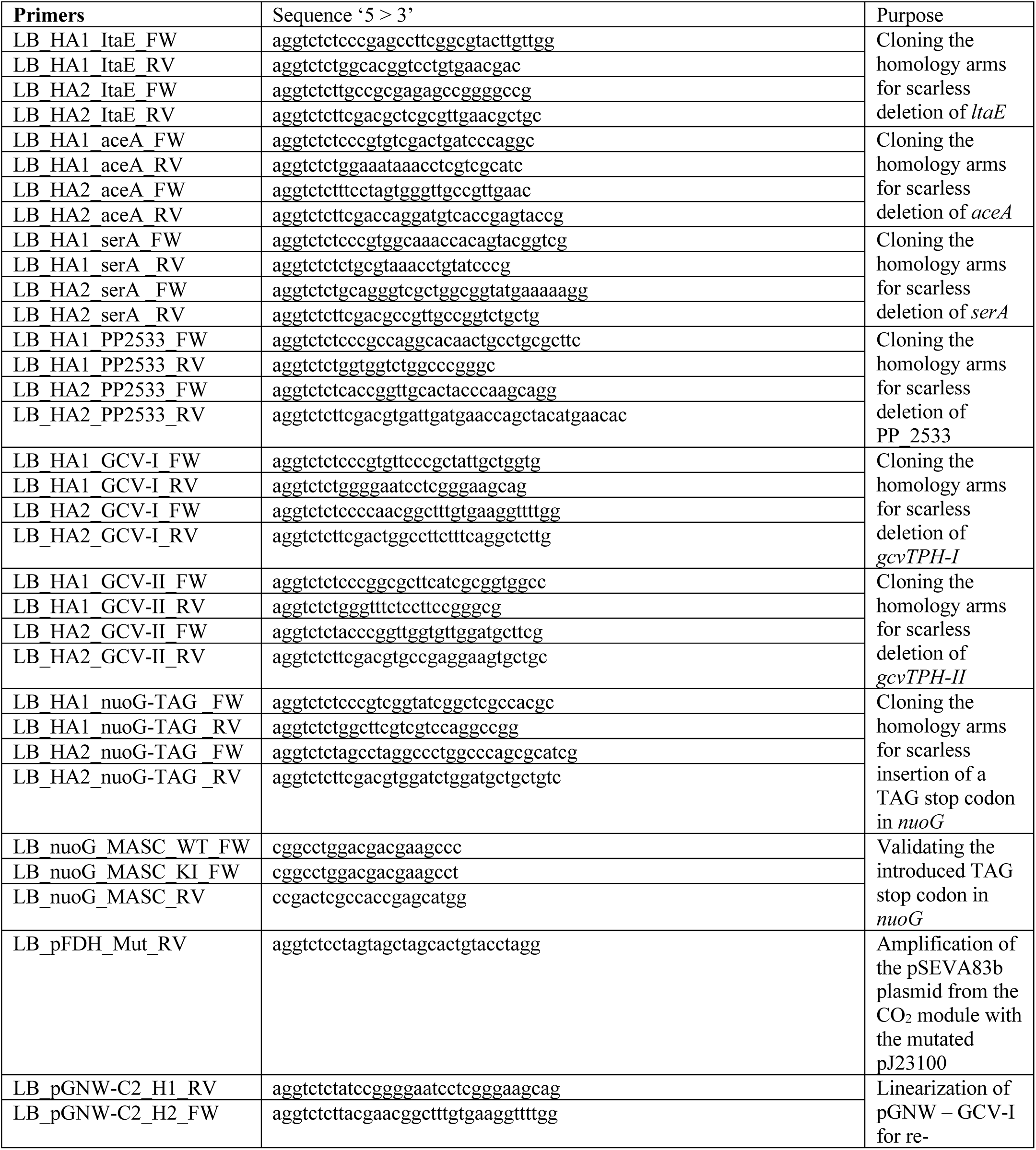
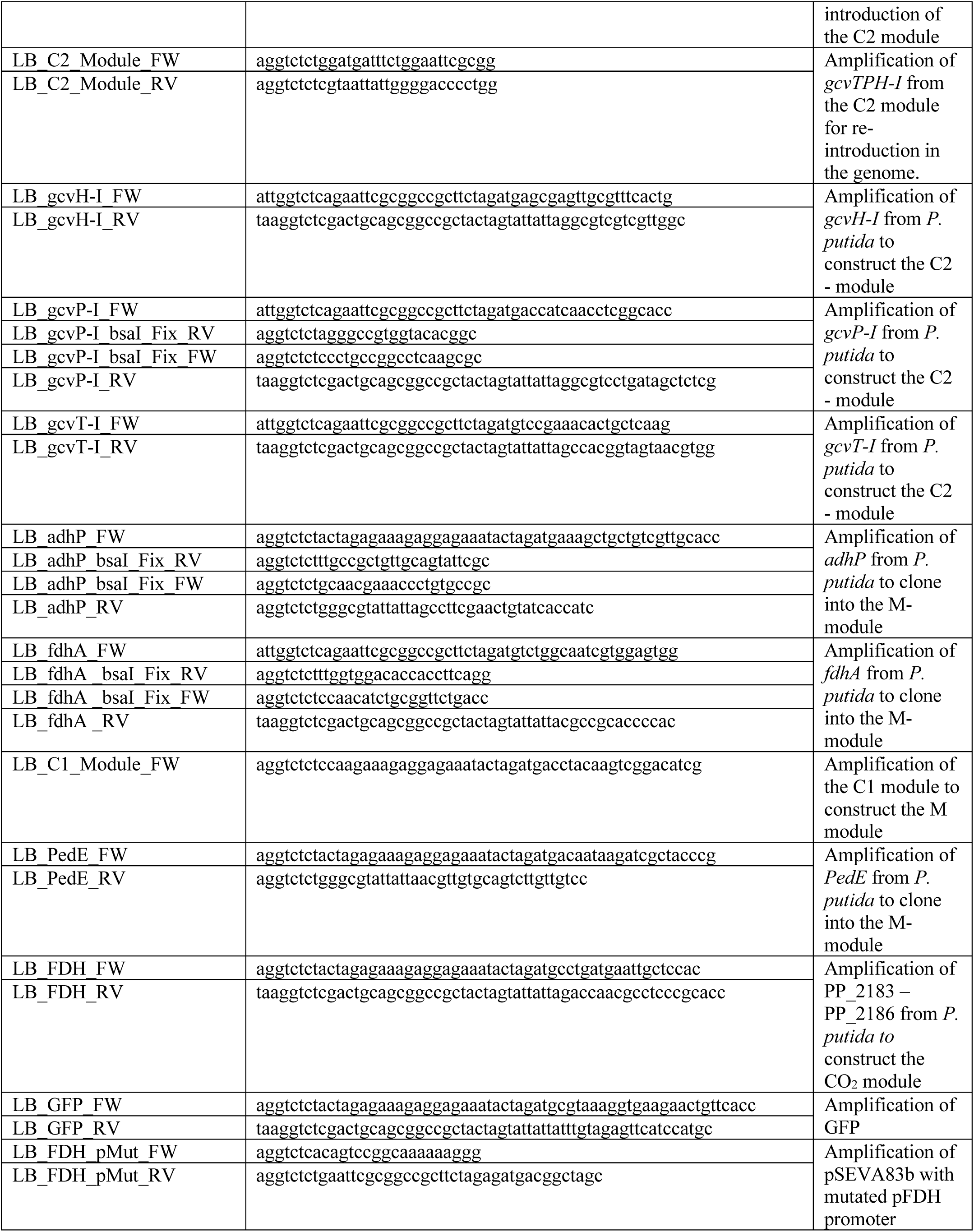
Primers used in the present study.

**Table S 3.** Mutations identified in C1 -S –Evo after short-term evolution.

| Genome position | Gene name (Locus ID) | Mutation | Annotation | Gene annotation |
| --- | --- | --- | --- | --- |
| 9126 | *rpmH* ← / → *dnaA* | +G | intergenic (-180/-416) | N.A. |
| 278744 | PP_0224 ← / ← *sctC* | +A | intergenic (-239/+75) | N.A. |
| 2472058 | PP_2166 | Δ1 bp | coding (445/483 nt) | Anti-anti-sigma factor |
| 4662562 | *nuoG* (PP_4124) | C→T | Q476* (CAG→TAG) | NADH-quinone oxidoreductase subunit G |

**Table S 4.**
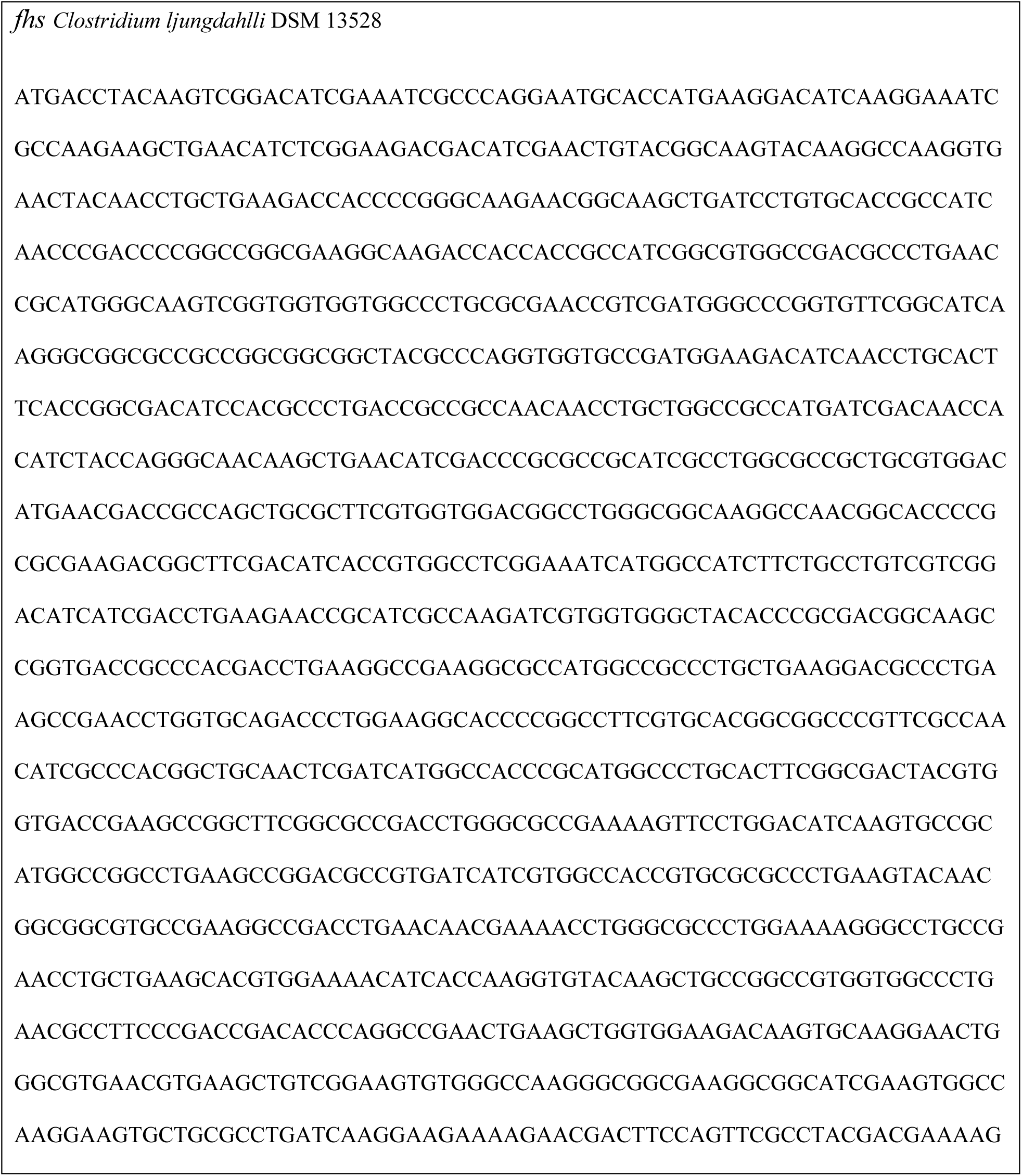
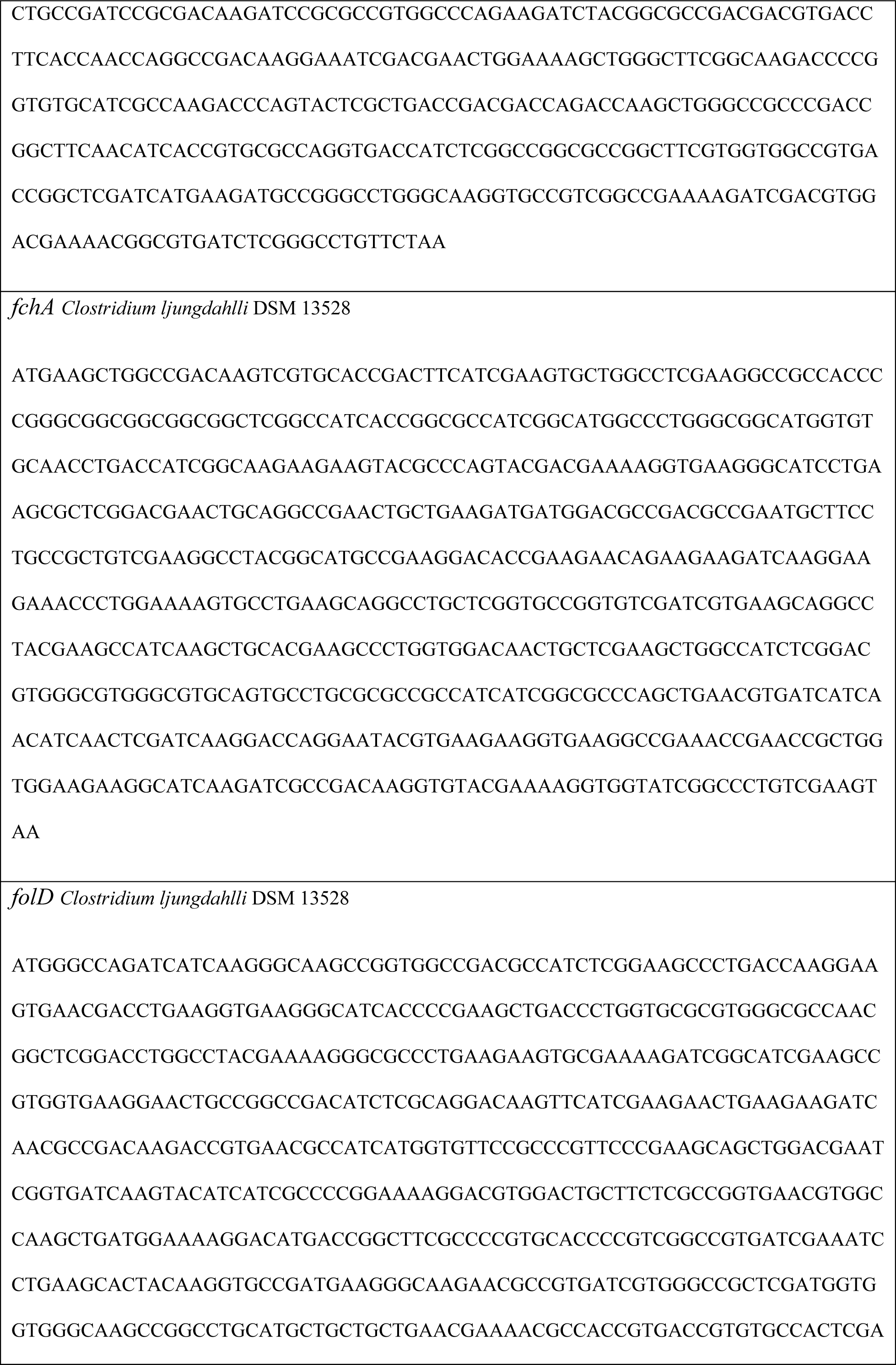
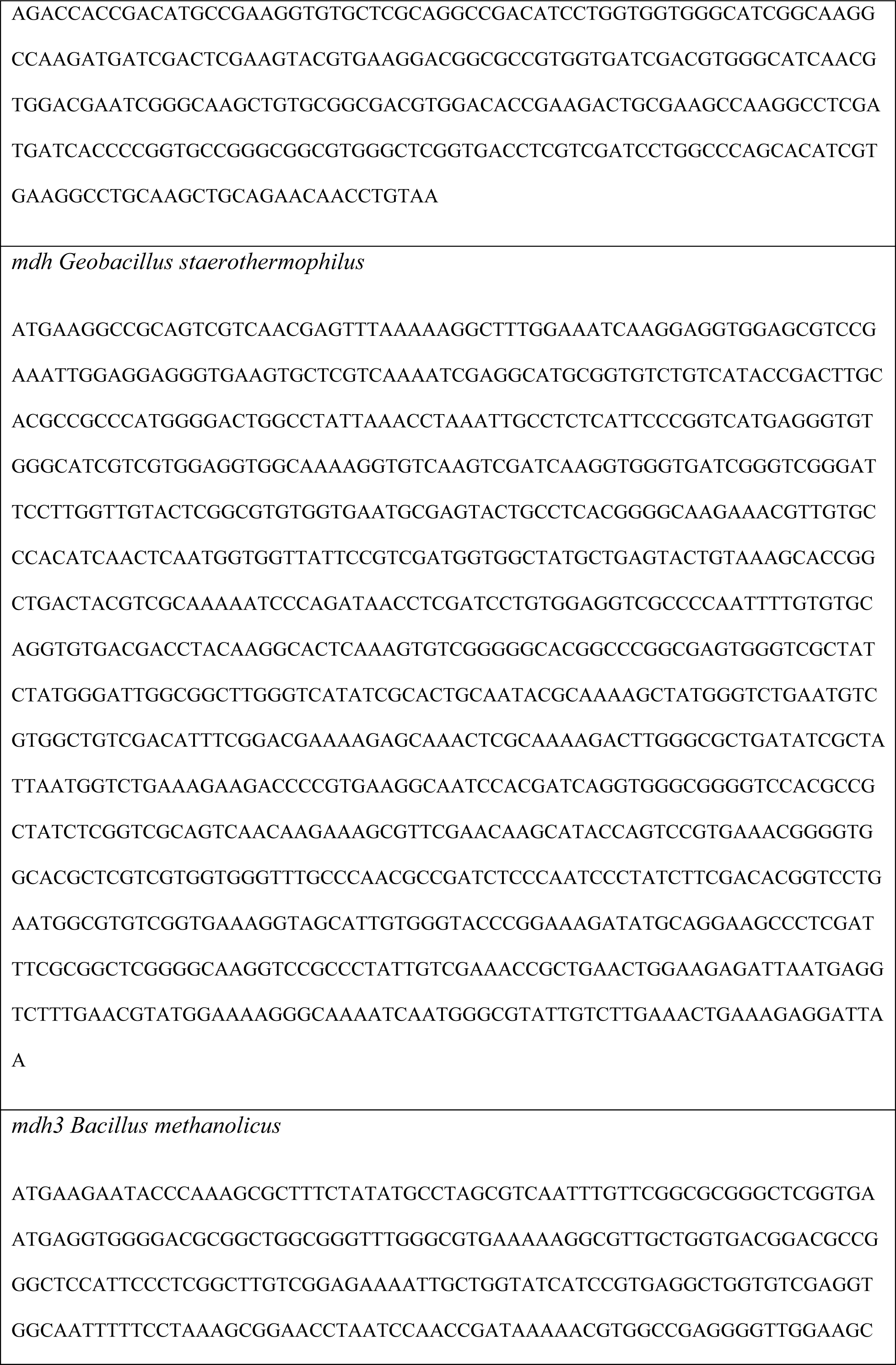
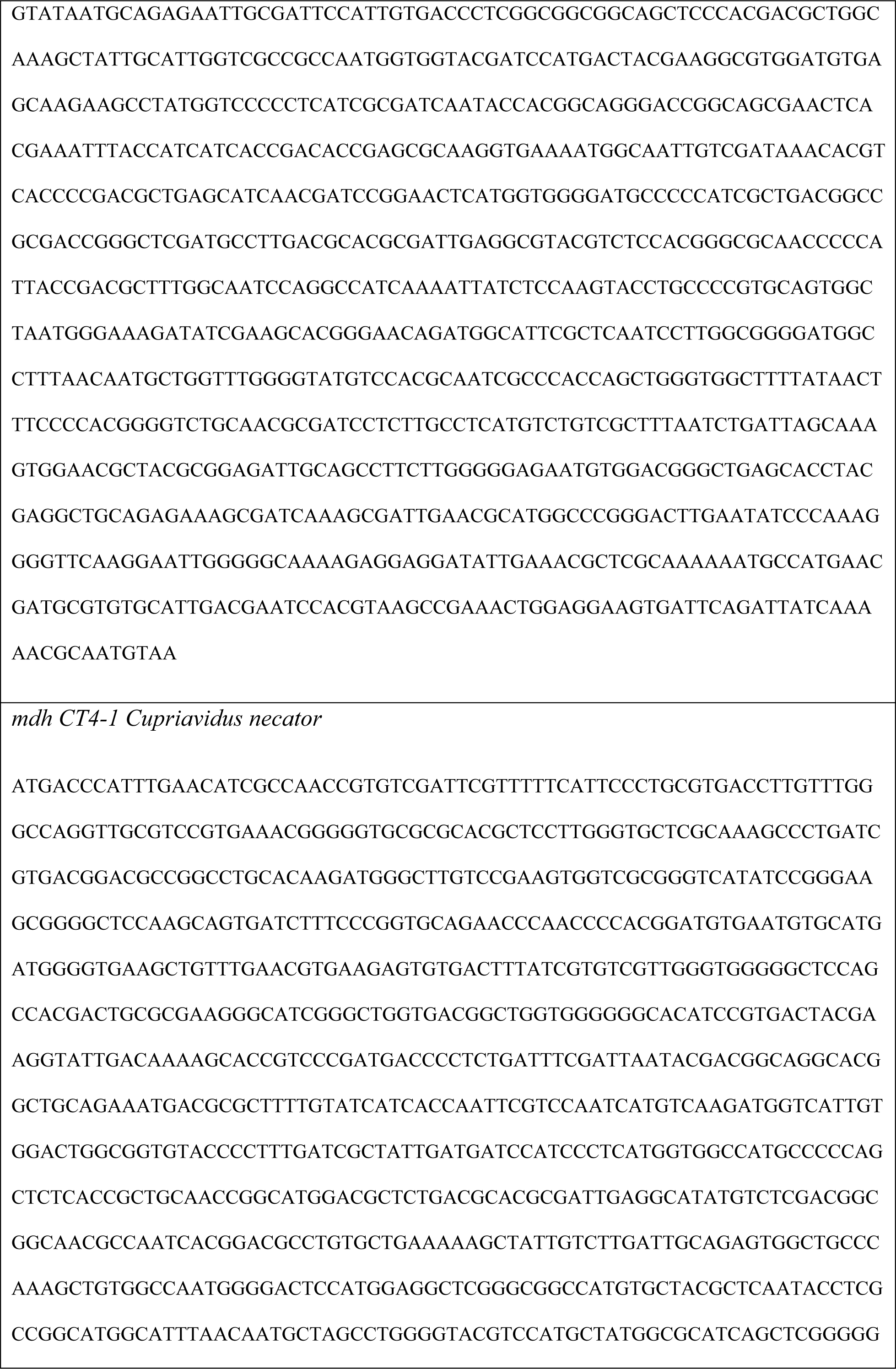
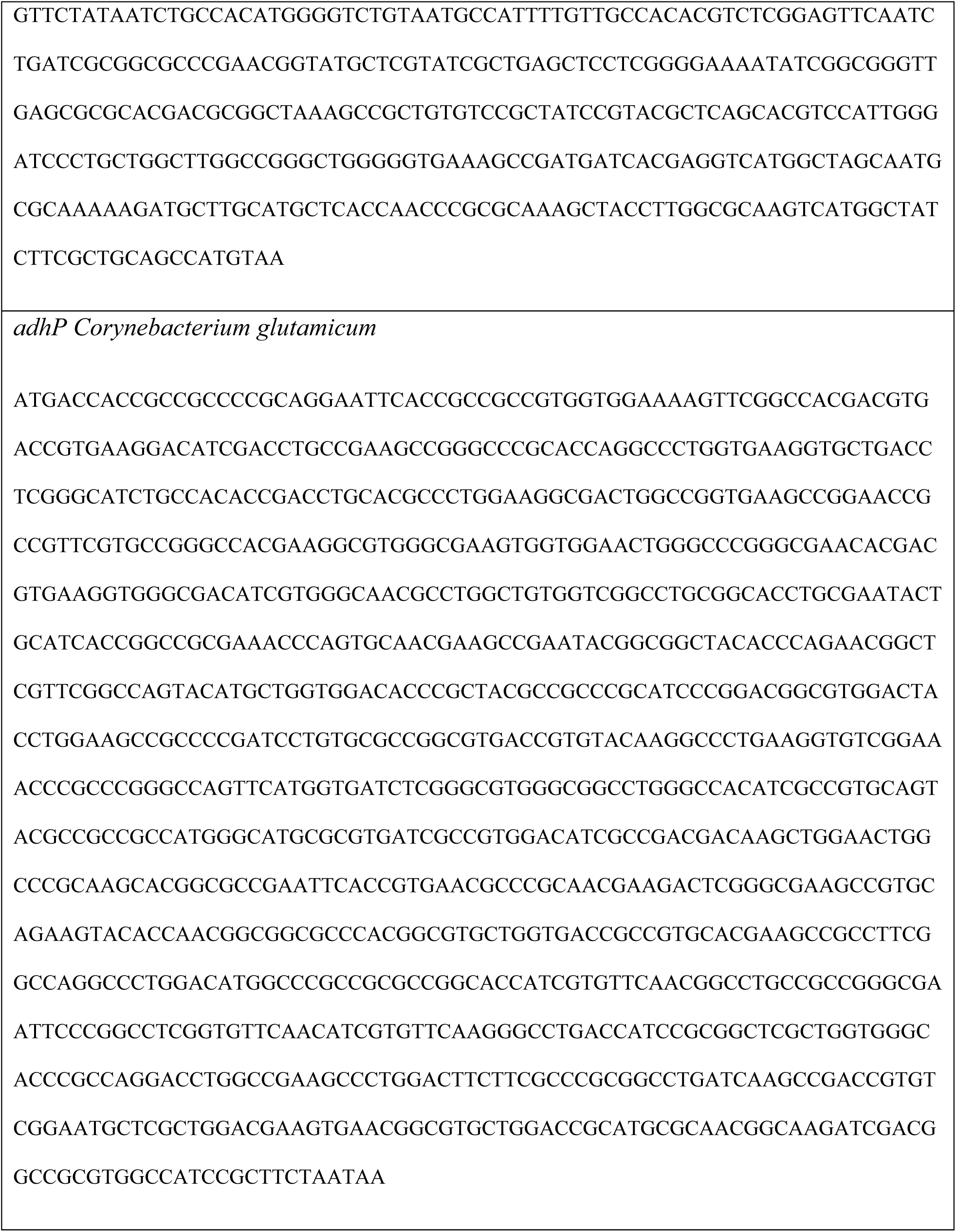
Codon optimized genes for the C1 module and M module.

